# Deciphering the species-level structure of topologically associating domains

**DOI:** 10.1101/2021.10.28.466333

**Authors:** Rohit Singh, Bonnie Berger

## Abstract

Chromosome conformation capture technologies such as Hi-C have revealed a rich hierarchical structure of chromatin, with topologically associating domains (TADs) as a key organizational unit, but experimentally reported TAD architectures, currently determined separately for each cell type, are lacking for many cell/tissue types. A solution to address this issue is to integrate existing epigenetic data across cells and tissue types to develop a species-level consensus map relating genes to TADs. Here, we introduce the *TAD Map*, a bag-of-genes representation that we use to infer, or “impute,” TAD architectures for those cells/tissues with limited Hi-C experimental data. The TAD Map enables a systematic analysis of gene coexpression induced by chromatin structure. By overlaying transcriptional data from hundreds of bulk and single-cell assays onto the TAD Map, we assess gene coexpression in TADs and find that expressed genes cluster into fewer TADs than would be expected by chance, and show that time-course and RNA velocity studies further reveal this clustering to be strongest in the early stages of cell differentiation; it is also strong in tumor cells. We provide a probabilistic model to summarize any scRNA-seq transcriptome in terms of its TAD activation profile, which we term a TAD signature, and demonstrate its value for cell type inference, cell fate prediction, and multimodal synthesis. More broadly, our work indicates that the TAD Map’s comprehensive, quantitative integration of chromatin structure and scRNA-seq data should play a key role in epigenetic and transcriptomic analyses.

*Software availability*: https://tadmap.csail.mit.edu

**Graphical Abstract:** 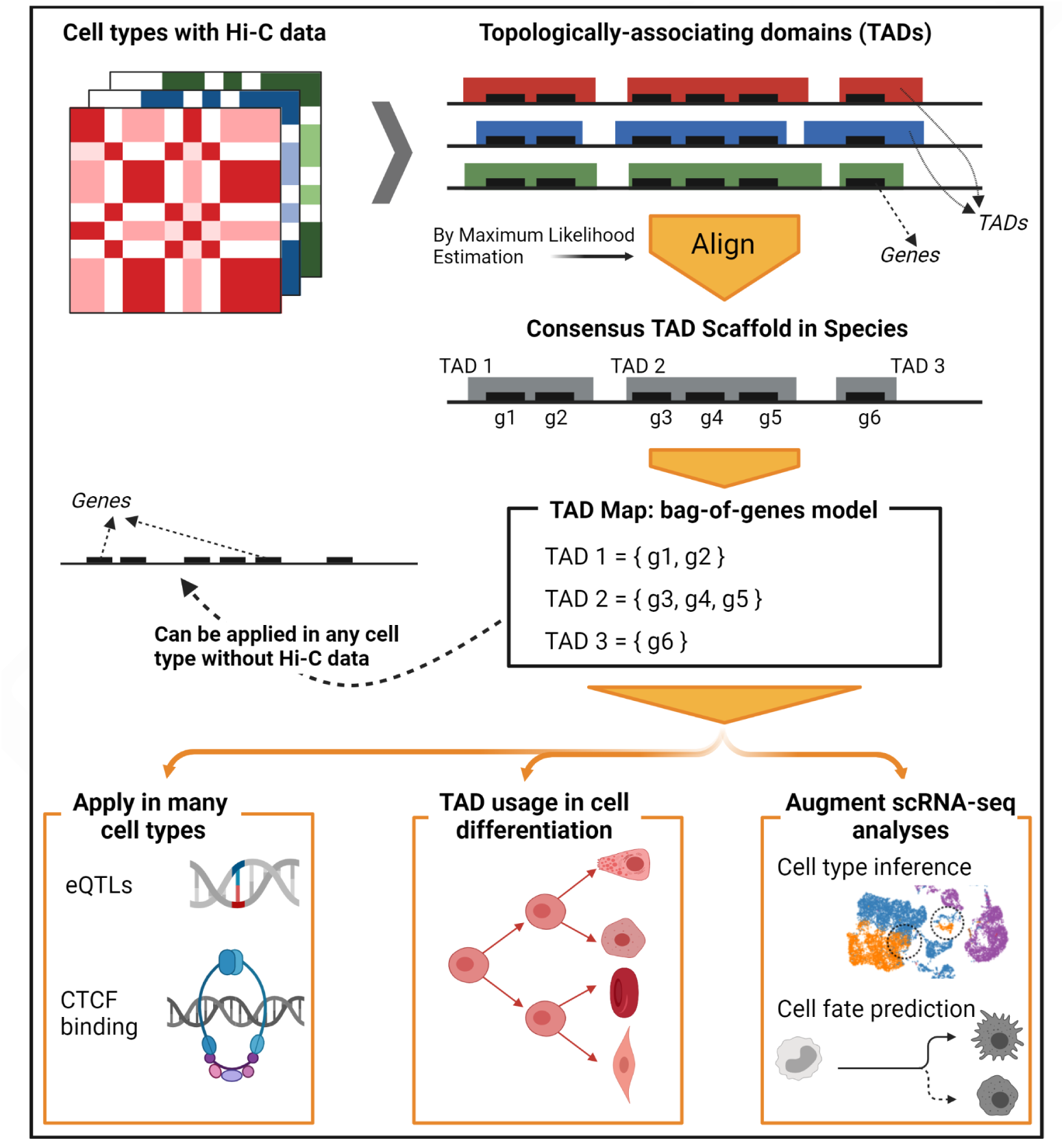

## Introduction

Advances in chromosome conformation capture technologies such as Hi-C, GAM, SPRITE, and ChIA-PET [1, 2] have informed how the 3-dimensional (3D) structure of the genome regulates gene expression, and in particular the role of topologically associating domains (TADs) in governing gene regulation. TADs, contiguous stretches of the genome where chromatin frequently self-interacts, represent a key level in the hierarchical organization of chromatin’s 3D structure [3, 4]. In cell types where high-throughput chromosome conformation capture (Hi-C) data is available [5–7] and TADs have been estimated, they have been shown to play an important role in regulation of gene expression [8–11]. Symmons et al. [12, 13] showed that at the *Shh* gene locus, enhancer activity within the encompassing TAD was pervasive and not substantially dependent on genomic distance . More broadly, Long et al. [14] discussed the potential role for TADs as bounding regions that bring genes in close proximity to their enhancers. Disruption of TAD boundaries has been linked to severe changes in gene expression and also implicated in disease [15–18]. A systematic and quantitative characterization of the link between TAD architecture and transcription— one that would hold across cell types and differentiation stages— could offer fundamental insights into cellular regulation.

Unfortunately, our current understanding of the connection between TAD architecture and transcription is limited. While cell differentiation studies of bulk and single-cell RNA-seq (scRNA-seq) data have documented a systematic loss of transcriptional diversity that marks cell differentiation [19], the corresponding systematic changes in a cell’s chromatin structure associated with this transcriptional change remain poorly understood. Other work on TADs has been limited to investigating gene expression in a specific genomic region [13, 20] or a limited set of cell types [11, 21, 22], or has instead focused on a mechanistic understanding of the molecular processes at work [3, 23]. Recently, Mateo et al. [24] described a multimodal single-cell technique to assay both DNA structure and gene expression but the study was limited to the *Hox* gene cluster in *Drosophila*. Also, though many prior studies broadly concur on the importance and role of TADs, some do not [25, 26]. Unfortunately, studies that could help separate general principles from one-off phenomena have been limited by experimentally-determined TAD architecture’s availability for only some cell types and differentiation conditions, far fewer than for which RNA-seq data is available. Especially in a single-cell context, the large number of potential chromatin contacts each of which is observable only twice per diploid cell suggest that acquiring robust Hi-C data to accompany a scRNA-seq study will remain a challenge. Instead, analogous to genotype imputation for genome-wide association studies [27], we seek to impute a TAD architecture broadly compatible with any RNA-seq study, obviating the need for study-specific Hi-C data.

Here, we construct the “TAD Map,” a representation of TADs (and an algorithm for computing it) that abstracts away details of minor chromatin structural variations and is strongly conserved across cell types, thus making it useful for species-level investigations (**Figure 1**). While chromatin conformation does vary at the cellular level, the overall architecture of TAD boundaries has nonetheless been found to be broadly consistent across cell types [8, 11, 28–30]. We therefore propose a *bag-of-genes* model for TADs, characterizing them in terms of the genes they partially or fully contain and find that in this representation, TAD architectures become even more consistent across cell types. We infer the species-level TAD architectures for human and mouse from Hi-C data [7, 21], applying a maximum likelihood estimation procedure. We show that the genome segmentation implied by the TAD Map is broadly informative, demonstrating its agreement with gene coexpression, expression quantitative trait loci (eQTL) localization, and CCCTC-binding factor (CTCF) binding sites in a variety of cell and tissue types and across diverse transcriptional contexts.

**Figure 1:**
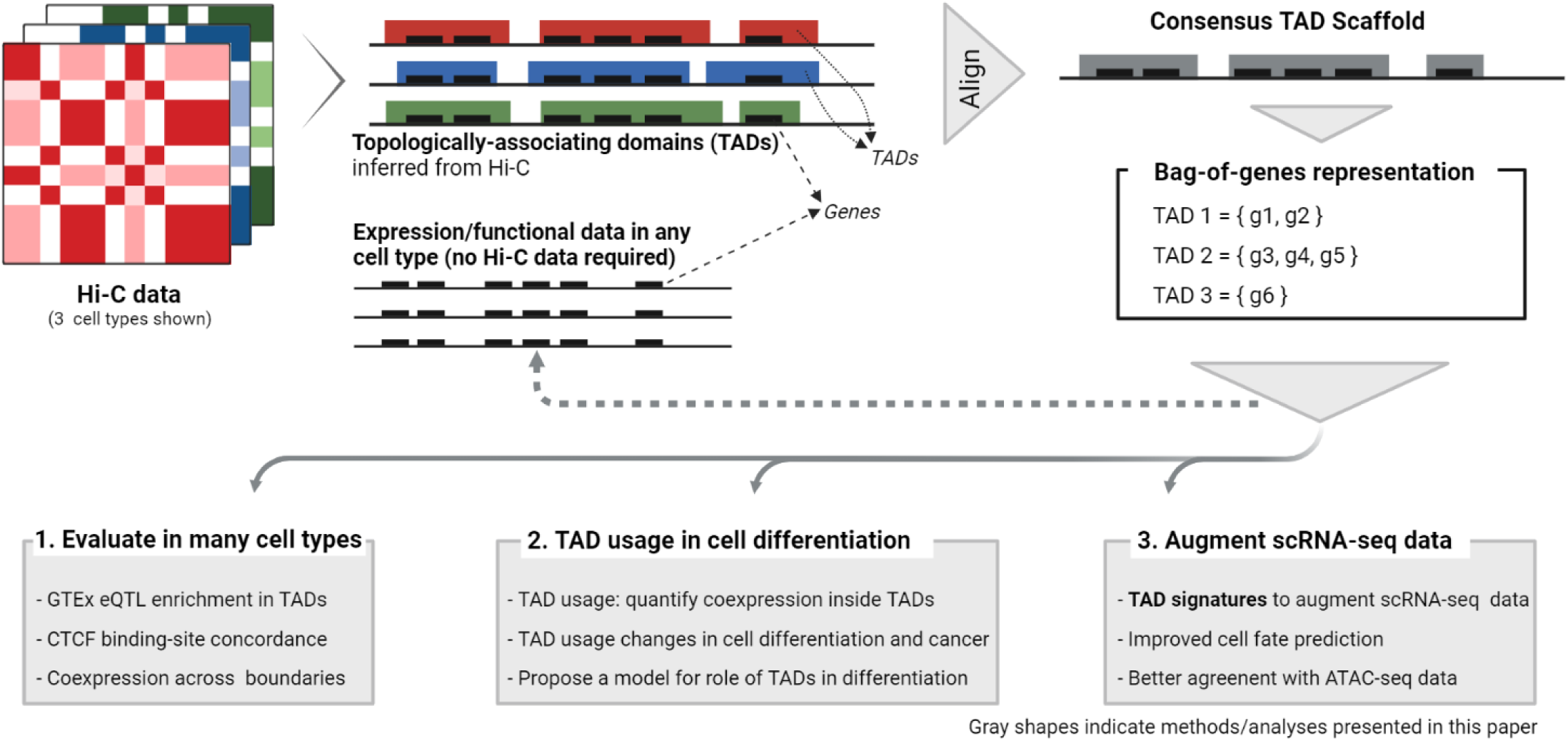
**Overview**: Topologically associating domains (TADs) are a key folding unit of the chromatin. Currently inferred only from cell-type specific Hi-C data, TAD architectures have been observed to have good agreement across cell types. We find this agreement to be even stronger when TADs are represented simply as sets of genes, suggesting a bag-of-genes representation would have species-wide applicability. Applying maximum likelihood estimation, we compute a consensus TAD scaffold from Hi-C data and the corresponding bag-of-genes representation (the *TAD Map*). The TAD Map enables us to “impute” the high-level chromatin structure in each cell/tissue. (i) We demonstrate that the genome partitioning the TAD scaffold implies agrees with functional genomic data across a variety of cell/tissue types. (ii) The TAD Map further uncovers the role of TADs in the transcriptional changes that mark cell differentiation. (iii) We introduce *TAD signatures*, a probabilistic model of TAD activation inferred from single-cell RNA-seq (scRNA-seq) readouts, showing how they facilitate greater accuracy and robustness in downstream scRNA-seq analyses (e.g., cell fate prediction).

The key conceptual breakthrough enabled by the TAD Map is to relate the transcriptional profile of any cell/tissue sample to the genome’s overall chromatin structure without the need for direct Hi-C data on that sample. Across hundreds of bulk RNA-seq, scRNA-seq, and RNA-velocity datasets, we investigated how the gene groupings implied by the TAD Map relate to gene coexpression. We found that gene expression is clustered into TADs, with the extent of this clustering changing during cell differentiation and being most pronounced in early stage, stem-like cells. Concordantly, we also found tumor cells to display substantially greater clustering of expression than normal cells. Using RNA velocity data to investigate the TAD localization of newly (de)activating genes revealed that the loss of transcriptional diversity during cell differentiation acts non-uniformly on TADs: newly active genes are more likely to hail from hitherto inactive TADs rather than from partially active TADs. Seeking additional evidence that TADs may provide an environment where adjacent genes are co-transcribed, we categorized adjacent gene pairs by whether they had the same or opposing orientations. Analyzing coexpression, the frequency of erroneous intergenic transcripts, and synteny from human to mouse, we found that shared orientation of adjacent genes is statistically significant inside TADs but not outside them.

To incorporate the TAD Map into standard scRNA-seq analyses such as cell type inference, cell fate prediction, and multimodal synthesis, we introduce the concept of a cell’s *TAD signature*, a decomposition of its gene expression profile into per-TAD activity estimates. We design a probabilistic generative model to infer TAD signatures from any scRNA-seq dataset, finding that the aggregation of transcriptional activity by TADs exposes additional biological signal not directly accessible from raw gene expression data. We apply TAD signatures to improve cell type inference, better distinguishing between mature and progenitor luminal cells in breast tissue, and also use them to identify the most informative genes for cell fate prediction.

TAD Map’s incorporation of epigenetics into analyses of transcriptional observations facilitates the application of insights from the genome’s 3D structure to inform analyses of bulk and single-cell RNA-seq (scRNA-seq) data that are now the bedrock of many biological investigations.

## Results

### Generating the consensus TAD Maps for human and mouse

It has been previously noted that TAD positions show substantial overlap across cell types [8, 11, 28, 29]. Extrapolating from this observation, we hypothesized that these cell type-specific architectures are variations of a consensus TAD scaffold that is applicable species-wide. To test this hypothesis, we first quantified the extent to which TAD architectures called for different cell types in a species are mutually consistent. Sourcing all TAD definitions in the TADKB database [31], we investigated TAD architectures for 7 human and 4 mouse cell types called using the Directionality Index technique [8]; the use of TADKB harmonized the calling of TADs on data from varying sources (**Methods**). We found substantial overlap between the TAD architectures: across TAD architectures for the 7 human cell types, 92.6% of TAD boundaries had a near-exact match (within 50 kb) in at least one other cell type’s architecture. However, the overlap was not complete: only 31.7% had a near-exact match in *all* the other cell types. The corresponding scores for the 4 mouse cell types were 69.9% and 13.4%, respectively. **Figure 2A** illustrates the overlaps on example segments from the human and mouse genomes.

**Figure 2:**
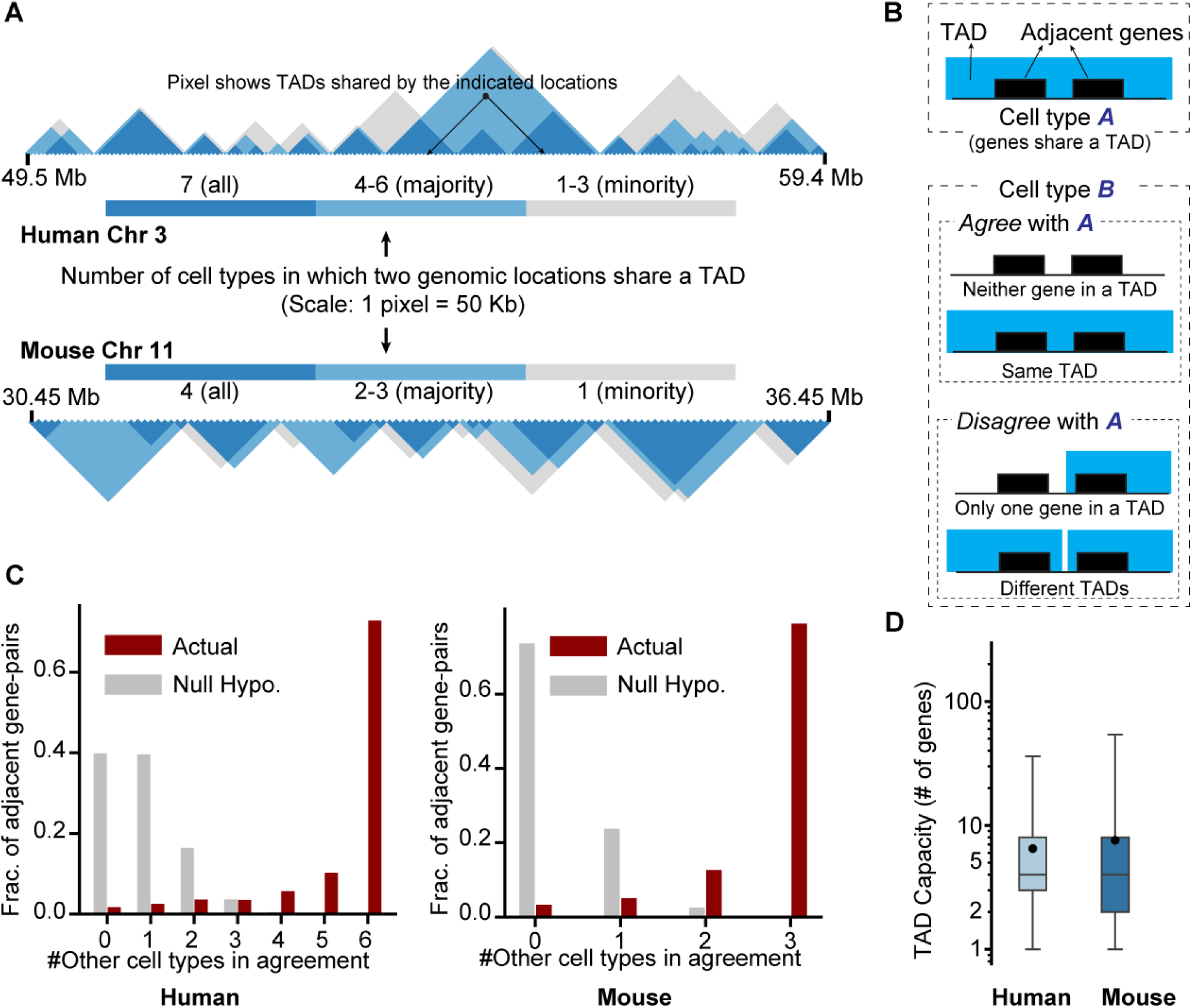
**Agreement between cell-type specific TAD architectures and the constructed consensus TAD scaffold**: **A**) On representative segments of the human and mouse genomes, overlap of cell-type specific TAD architectures (7 cell types for human, 4 four mouse) are shown. Genomic positional range (in Mb) is shown on the horizontal axis and the number of cell types in which a pair of genomic loci co-occupy a TAD is indicated by the shaded triangles. In each species, the majority of the cell-type specific TAD architectures are in agreement on most genomic loci pairs; in many cases, *all* TAD architectures are in agreement. **B**) Schematic for the statistical test to evaluate if two bag-of-genes TAD representations are identical: given an adjacently-located gene pair that shares a TAD in at least one cell type’s TAD architecture, we mark other cell type(s) to be in agreement if the gene pair is not separated by a TAD boundary. **C**) For each adjacent gene pair that shares a TAD in at least one cell type, the number of other cell types where the genes share a TAD. Under this test, 72.7% of adjacent gene pairs in human (78.9% in mouse) show complete agreement across all cell-type specific TAD architectures. The expected count under a simple null hypothesis (Methods) is below 1 in both species. **D**) Our consensus TAD scaffolds for each of human and mouse, computed by maximum likelihood estimation, yield 3036 (3181) TADs with an average length of 879 kb (845 kb) in the human (mouse) genome, with the median TAD in both species containing 4 genes. In this figure, the box represents the 25-75*^th^*percentile range, and the whiskers represent 1-99*^th^*percentile range. Human and mouse TAD scaffolds correspond well with each other, though we found the mouse scaffold to contain more large-capacity TADs.

Reasoning that a lower-resolution representation of TADs could smooth these boundary variations, we abstracted a TAD simply as a bag (i.e., set) of protein-coding genes partially or fully contained in it. The key advantage of this representation (the “TAD Map”), which is inspired by the bag-of-words model common in natural language processing tasks, is that any TAD boundary variations that do not change the gene set become irrelevant. This representation of TADs still retains sufficient detail for many analyses (e.g., relating gene coexpression with TAD-level chromatin structure), though it does lose some details (e.g., locations of enhancers).

We found the bag-of-genes representation of TADs to be remarkably consistent across cell types. To compare two bag-of-genes representations, it is sufficient to consider gene pairs that are adjacent on the genome; since TADs are arranged sequentially along the genome, the representations will be identical if they agree on the grouping of each such pair. For each adjacent gene pair that co-occupies a TAD in some cell type, we checked if the TAD representations in other cell types preserved this grouping (Figure 2B). For 72.8% (78.9%) of such gene pairs in human (mouse), representations for *all* the remaining cell types were in agreement; for 92.3% (91.6%) of the pairs, the majority of them were in agreement. As an alternative quantification, we compared these estimates to a null hypothesis where TAD boundaries are independently dispersed between gene boundaries in each cell type while preserving the number of TADs. As **Figure 2C** shows, for a gene pair that co-occupies a TAD in one of the architectures, the expected number of other cell-type architectures that agree with this grouping under the null hypothesis is just 0.85 (human) and 0.29 (mouse), implying that extent of agreement actually seen is highly significant (one-sided binomial test, *p* = 7.3 × 10^−75^ for human, 1.7 × 10^−4^ for mouse).

Leveraging this observation, we constructed species-wide TAD “scaffolds” for mouse and human. Serving as a precursor to the TAD Map, the scaffold demarcates consensus estimates of TAD boundaries. We compute it from cell type-specific data by maximum likelihood estimation: partitioning a chromosome into 50 kb segments, for every pairwise combination of segments we score the aggregate experimental evidence that the segments share a TAD. We then search for the optimal set of non-overlapping genomic intervals maximally supported by the evidence, noting that certain genomic regions might not be covered by any TAD (**Methods**). We solve the optimization with a dynamic programming algorithm, building a consensus TAD scaffold for human with 3036 TADs (mean length 879 kb, with the median TAD containing 4 protein-coding genes) and a mouse TAD scaffold with 3181 TADs (mean length 845 kb, median 4 protein-coding genes per TAD) (**Figure 2D**). From this consensus TAD scaffold, we inferred the bag-of-genes representation (i.e., the TAD Map), associating each TAD with the set of genes fully or partially contained in it, with genes that spanned two TADs being associated with both (**Discussion**).

### The TAD scaffold agrees with functional genomic data on a variety of cell types

We evaluated the TAD scaffold’s concordance with functional genomic data on a wide cross-section of cell, tissue and organ types, reasoning that if the genome partitioning it implies is accurate in diverse contexts, strong within-TAD functional associations should be observed correspondingly. In our evaluation, we considered two measures of functional association: expression quantitative trait loci (eQTL) and coexpression. Additionally, to assess the mechanistic evidence for the TAD scaffold we also computed its agreement with CTCF ChIP-seq data in a variety of tissue types.

#### Expression quantitative trait loci

By relating a gene’s expression to genotypic variability around its genomic locus, eQTLs can indirectly indicate the genomic range over which epigenetic control of gene expression is likely exercised. We evaluated human TAD scaffold predictions on the entire compendium of single-tissue eQTL data from the Genotype-Tissue Expression Portal (GTEx) v8 [32], with 49 datasets covering a variety of organs and tissues. Aggregating across these and limiting the analysis to eQTLs with p-value less than 10^−5^, we estimated the probability *p* (*d*) of a genomic locus at distance *d* from a gene’s transcription start site (TSS) being associated with an eQTL for the gene (**Methods**); we averaged this estimate over TSS-eQTL pairs that co-occupy a TAD and separately over pairs that do not. As has been observed previously, the probability *p* (*d*) declines with *d* [33, 34]. However, we found the rate of decline to be significantly lower inside TADs (**Figure 3A,B**). Except for eQTLs very close to the TSS (*d <* 30 kb), the probability *p* (*d*) is significantly higher inside TADs (**Figure 3A,B**), with a median fold change in *p* of 1.27 (upstream eQTLs) and 1.37 (downstream eQTLs) when comparing TSS-eQTL pairs that share a TAD to those that don’t (*p*-value = 4.8 × 10^4^ and 4 × 10^−4^, respectively; Wilcoxon rank sum test). We limited the upper bound of our evaluation range to 1 Mb, since the GTEx corpus limits single-tissue eQTL reports to this range.

**Figure 3:**
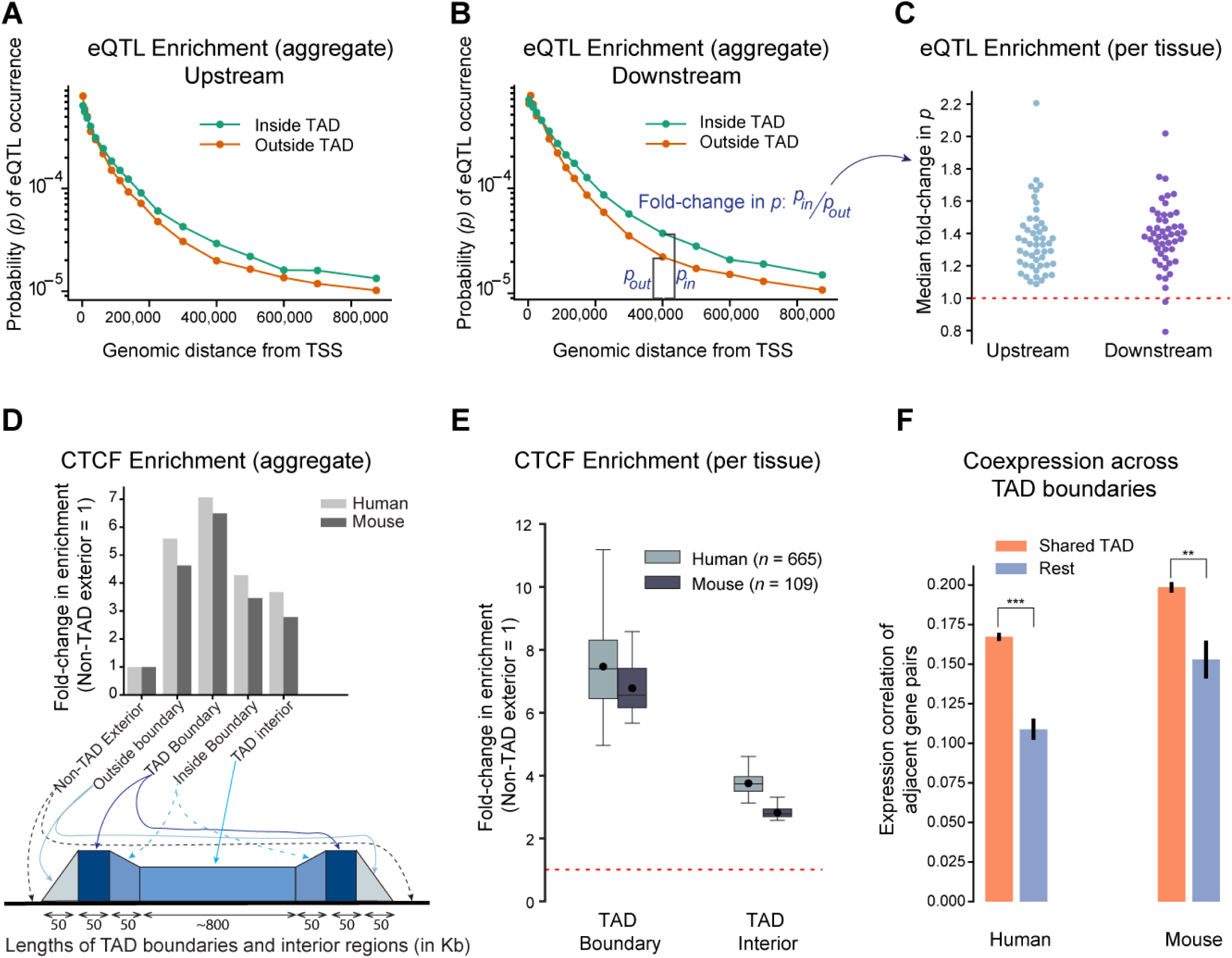
**The TAD scaffold is functionally meaningful across a variety of tissue and cell types: A, B, C**) To assess the TAD scaffold’s functional informativeness, we compared expression quantitative trait loci (eQTL) prevalence inside and outside TADs. Across all single-tissue eQTLs from GTEx v8, we plotted the average per-gene frequency of eQTL occurrence against genomic distance from the gene’s transcription start site (TSS), categorizing the eQTL by whether its locus and the gene TSS share a TAD. A,B) Aggregated across tissue types, genomic loci that shared a TAD with the gene’s TSS were associated with a higher eQTL frequency (*p*= 0.0001 for both upstream and downstream loci, one-sided Wilcoxon rank sum test). (C) This was also true individually for most tissues/organs, indicating the TAD scaffold’s informativeness across many tissue types. **D, E**) Since CCCTC-binding factor (CTCF) is thought to be part of the mechanism for TAD formation in the chromatin, we correlated CTCF binding activity in a variety of tissue and cell types with the predicted consensus TAD scaffold boundaries (produced at a 50 kb resolution) . Across 665 human and 109 mouse CTCF ChIP-seq assays (ENCODE), there was significant CTCF binding enrichment at TAD boundaries and also within TADs. CTCF enrichment falls off at either side of our predicted TAD boundaries, suggesting the predictions are approximately correct. E) In further support of the TAD scaffold’s applicability to diverse cell/tissue types, enrichment of CTCF binding at TAD boundaries was seen at the level of individual tissues (Bonferroni-corrected *p* < 10^−50^ in all cases, one-sided binomial test). In this figure, the box represents the 25-75*^th^*percentile range, and the whiskers represent 1-99*^th^*percentile range. **F**) Investigating gene pairs located adjacently along the genome, we find significantly higher coexpression between pairs that share a TAD (*p* = 2.1 × 10^−8^ in human and 0.035 in mouse, one-sided t-test; evaluated over 863 bulk RNA-seq datasets from ENCODE)

Notably, though the 49 GTEx datasets correspond to a diversity of tissue and organ types (e.g., blood, brain, liver, skin, etc.), we observed a higher in-TAD eQTL occurrence rate not just overall but in almost *all* of them individually (**Figure 3C**). Thus, despite being estimated from data on a limited set of cell types, the consensus TAD scaffold can delineate regions of enhanced functional association in a variety of tissue and cell types.

#### Agreement with ChIP-seq

We also evaluated the TAD scaffold’s agreement with the mechanistic model of chromatin folding. A key mechanism for TAD formation is thought to be CTCF-mediated loop extrusion, with cell type-specific studies of TAD formation finding CTCF binding to be enriched at TAD boundaries [35, 36]. There may also be higher CTCF binding activity inside TADs, helping shape the chromatin’s hierarchical organization [3]. Reasoning that our consensus TAD scaffold should be similarly supported by CTCF binding evidence, we acquired CTCF ChIP-seq data from the ENCODE database [37], consisting of 281 human and 28 mouse studies (comprising 665 and 109 replicates, respectively). These assays again covered a variety of cell types and disease states, allowing us to test if the TAD scaffold generalizes well species-wide.

Using the rate of CTCF binding in non-TAD genomic regions as the baseline, we quantified the prevalence of CTCF ChIP-seq peaks inside TADs and at their boundaries (**Methods**). In both human and mouse, we found our results to confirm what has been previously observed: CTCF binding was most frequent at TAD boundaries and was substantially higher inside TADs than in the general genomic background (**Figure 3D**). Again, this was true not only in aggregate across all studies but also individually in a diverse set of cell types and disease states (**Figure 3E**). Next, seeking to assess whether our estimate of boundary locations is precisely correct or only approximately so, we also evaluated binding prevalence in 50 kb regions on either side of a TAD boundary (we recall that our TAD scaffold is inferred at a 50 kb resolution). Supporting the scaffold’s TAD boundary estimates, we found the regions adjoining the boundary to have substantially lower CTCF binding rates than seen at the TAD boundary itself. However, CTCF binding rate in these adjoining regions is still higher than the rate seen in the TAD exterior or interior. This could be explained by minor boundary variations across cell types (as is indeed known to happen) or if the “effective” TAD boundary is wider than our 50 kb definition. We note that the bag-of-genes model for TADs is designed to handle such ambiguity: as long as the gene memberships in a TAD are unchanged, minor variations in its boundaries are immaterial.

#### Coexpression of adjacent gene pairs

When evaluating the TAD scaffold, a possible null hypothesis is that the scaffold provides no additional information beyond the well-known fact that genes co-located on the genome are likely to have similar expression and function [38]. Under this null hypothesis, the genome partition demarcated by the consensus TAD scaffold is no more informative than any other partition. To test this hypothesis, we computed the coexpression of pairs of neighboring genes, grouping those pairs by whether or not they occupied the same TAD. Testing on 794 (human) and 69 (mouse) bulk RNA-seq datasets sourced from the ENCODE database (again encompassing a variety of cell types), we found that adjacent genes that share a consensus TAD display greater coexpression than adjacent gene-pairs that do not share a TAD (Pearson correlation 0.167 vs. 0.109, *p*=2.1 × 10^−8^ for human, and 0.198 vs. 0.153, *p*=0.035 for mouse; one-sided t-test after a Fisher’s r-to-z-transformation, **Figure 3F**).

### Gene transcription is marked by a selective usage of TADs

The TAD Map allows us to investigate if the chromatin structure of the genome implies a systematic bias towards certain transcriptional profiles over others. As noted previously, there already is isolated, qualitative evidence that TADs influence coexpression patterns by acting as units of epigenetic control. However, with the TAD Map we can now quantitatively investigate this connection across a diverse set of cell types and disease states, without any need for accompanying Hi-C data.

To measure the link between coexpression and TAD co-membership, we introduce a bootstrap-based statistical test. We designed this test because the correlation measure used for bulk RNA-seq data above is vulnerable to the noise and sparsity in scRNA-seq datasets (e.g., due to varying sequencing depths, platform effects, and dropouts). It is also difficult to adapt for more fine-grained analyses (e.g., distinguishing between highly vs. moderately active TADs). In contrast, our statistical test compares each transcriptome to bootstrap samples generated from the transcriptome itself, making it easier to aggregate estimates across disparate datasets. Specifically, the test compares a TAD’s observed “usage” against its expected usage: we define a TAD’s *usage* in the context of a given transcriptional profile, computing it as the number of genes in the TAD that are transcriptionally active.1 We assess transcriptional activity as a binary event, with a gene considered to be active if it has one or more transcripts (in a scRNA-seq transcriptome) or is in the top-5,000 genes by transcript count (in a bulk RNA-seq transcriptome). The test operates under the null hypothesis that activity of genes in the profile is independent of the TADs they occupy. For any transcriptional profile, we compare the observed TAD usages with what should be expected for the profile under the null hypothesis, estimating the latter from bootstrap samples generated by shuffling genes across TADs: the bootstrap samples vary in TAD usages while preserving TAD capacities and the total number of active genes (**Methods**). For any transcriptional profile, we then compare the observed count of TADs with at least one active gene (i.e., usage ≥ 1) to the expected count under the null hypothesis, expressing this in terms of a bootstrap z-score (**Figure 4A**, **Figure S1**). While the bootstrap samples are specific to a transcriptional profile, the conversion of observed values to a z-score allows us to standardize and aggregate our estimates across multiple datasets for greater statistical power.

**Figure 4:**
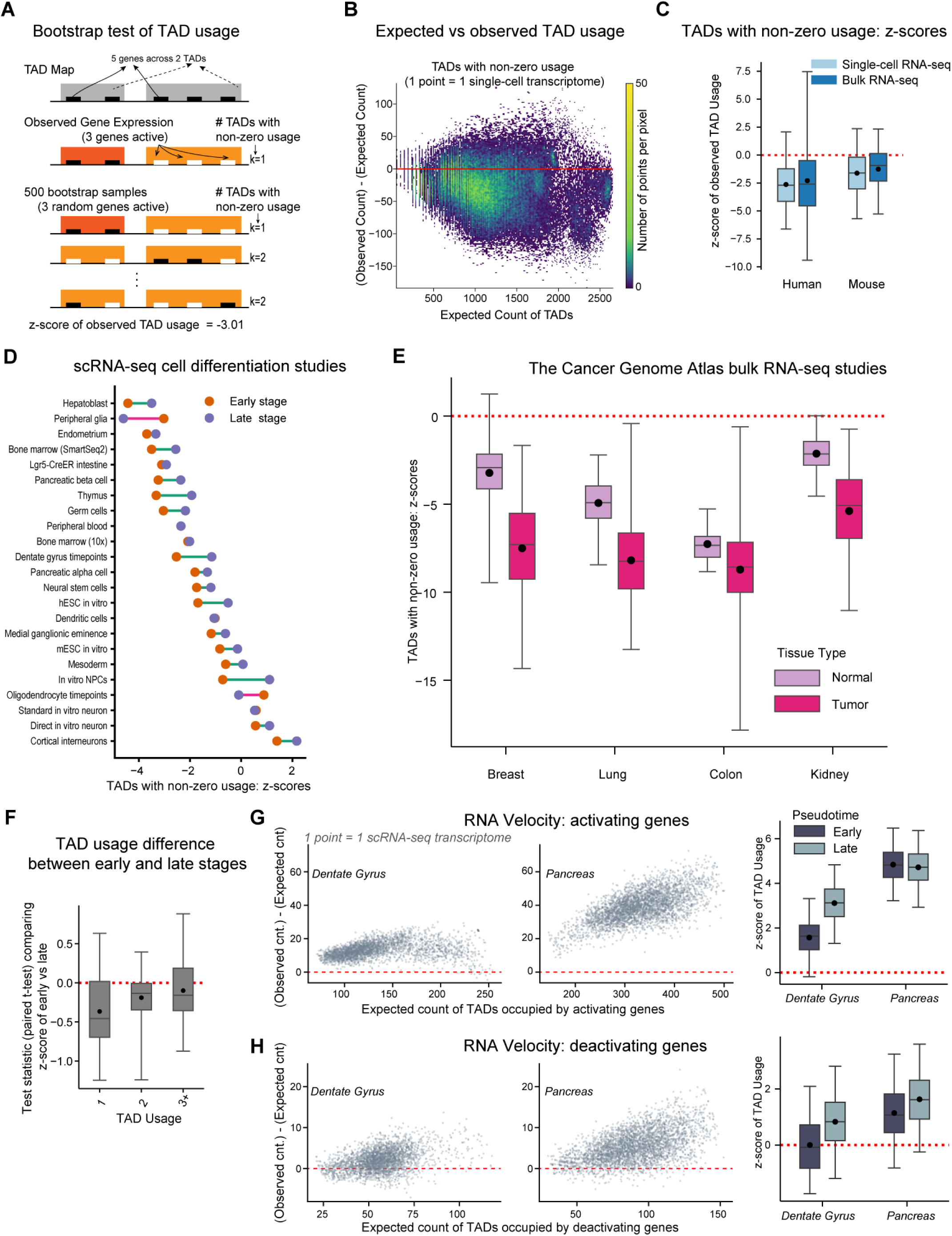
**(A-C) Gene expression is clustered into TADs and (D-H) this expression clustering is greater in both undifferentiated and cancer cells: A**) Bootstrap test to quantify the clustering of expressed genes into TADs by comparing against a null model with the same number of expressed genes, but randomly chosen from the entire set of genes. Each bootstrap sample randomizes genes across TADs, preserving TAD capacities and the total number of active/expressed genes. (**Figure S1** offers another example.) Converting each transcriptome’s observed TAD *usage* (i.e., number of genes in the TAD with non-zero expression) to a bootstrap z-score allows harmonization across disparate RNA-seq datasets, with negative z-scores indicating gene expression clustering into TADs. **B**) We analyzed 70,243 single-cell transcriptomes, counting the number of TADs with at least one expressed gene (i.e., TAD usage ≥ 1). Most cells demonstrate fewer TADs in use than expected, indicating clustering of active gene into TADs. **C**) Across these single-cell and 863 bulk transcriptomes in human and mouse, the number of TADs with usage ≥ 1 is significantly lower than expected (Bonferroni-corrected *p <* 10^−20^ in all cases, one-sided t-test). In all boxplots in this figure, the box represents the 25-75*^th^*percentile range, and the whiskers represent 1-99*^th^*percentile range. (D-H) Expression clustering is greater in undifferentiated and cancer cells: **D**) In each of 24 cell differentiation scRNA-seq studies with non-branching trajectories, cells were divided into early or late stage based on expert annotation [19]. Across most studies, early stage cells have fewer TADs with non-zero expression activity (i.e., usage ≥ 1), indicating a greater level of transcriptional clustering in these cells (Figure S2). **E**) Bulk RNA-seq data sets from The Cancer Genome Atlas (TCGA) corpus were analyzed for four cancers, with TAD usage in normal tissue compared to primary tumor. Like early stage cells, tumor cells also show greater expression clustering into TADs. **F-H**) Cell differentiation is marked by broad-based deactivation of previously active genes but also selective activation of lineage-specific genes. We find the latter is primarily responsible for the decrease of expression clustering in late stage cells. **F**) Comparing early- and late-stage cells in each scRNA-seq study, we evaluated the prevalence of TADs by usage levels, finding the largest difference in single-usage TADs, indicating late stage cells had more TADs with just one gene activated. **G,H**) Investigating two RNA velocity datasets (cell differentiation studies of the pancreas [47] and dentate gyrus [48], respectively), we mapped the loci of (de)activating genes into TADs. As measured by a bootstrap test, activating genes (top row) are likely to localize to hitherto unused TADs, consistent with the evidence from (F); the evidence for the clustering of deactivating genes (bottom row) is less clear. The box plots in F-H summarize the corresponding scatter plots by showing bootstrap z-scores; they additionally split the data into early and late stages as measured by *scVelo* pseudotime [46].

We applied this test to 70,243 scRNA-seq transcriptomes (aggregated from 30 datasets) and 863 bulk RNA-seq profiles (from ENCODE, as previously described); the datasets spanned two species (human and mouse), varying sequencing depths and library sizes, and seven distinct scRNA-seq protocols (the four most frequent were Fluidigm C1, 10x, Smart-seq2, and Drop-seq). The scRNA-seq datasets were curated by Gulati et al. [19], who sourced each dataset from a cell differentiation study; in the next section, we further analyze these datasets to investigate cell differentiation (**Methods**).

To evaluate if transcriptionally active genes cluster into a limited subset of TADs, we computed the number of TADs with non-zero activity (i.e., usage ≥ 1), reasoning that it should be lower than expected in the case of such clustering. We found this to be indeed the case: in 78.4% of the single-cell transcriptomes, the observed count of such TADs was less than expected (**Figure 4B**), with the population mean of the per-transcriptome z-score of this count being significantly negative (= -1.46, *p* ≈ 0, *t* statistic = -211.7, one-sided t-test).

To assess the robustness of this finding, we performed additional confirmatory analyses. We found the observation to hold true for both human and mouse data and for bulk and single-cell RNA-seq profiles (**Figure 4C**). Within single-cell RNA-seq datasets, we found it to hold on data from a variety of technologies (**Figure S2A**). Since our bootstrap test considers gene activation as a binary event, we wondered if our finding is an artifact of the binarization step. We chose the binarization approach since it lends itself to a robust and non-parametric test and avoids the need for a scRNA-seq transcript-count model that itself may be erroneous. To confirm that our findings were not an artifact of this binarization, we tested bulk RNA-seq data (where sequencing depths are greater than in scRNA-seq) and binarized it to select top-*k* genes, for various choices of *k*. We found the clustering to be robust to *k*, holding not only for highly-expressed genes but also for moderate and low expression (**Figure S2B**). Another indication of robustness is that high-usage TADs were more enriched than lower-usage ones: TADs with just a single active gene (usage = 1) were fewer than expected while TADs with 3 or more active genes (usage ≥ 3) were more frequent than expected (**Figure S2C**); we recall that the median consensus TAD contains four genes.

### Less-differentiated and cancer cells exhibit greater clustering of expression into TADs

A defining characteristic of both cell differentiation and tumorigenesis are broad changes in gene expression. Seeking to characterize how these transcriptional changes were influenced by the TAD architecture, we assessed how TAD usage patterns change during cell differentiation and between normal and tumor cells. For TAD usage in cancer, we analyzed bulk RNA-seq measurements of normal and primary-tumor tissue in blood, brain, lung, and renal cancers from The Cancer Genome Atlas database (TCGA, [39]). Performing the same statistical analysis as described previously, we found that TADs in which at least one gene was active (i.e., TAD usage ≥ 1) were relatively more prevalent in tumor cells (**Figure 4E**), indicating a greater clustering of transcriptionally-active genes into TADs. Since our statistical test adjusts for the number of expressed genes, this difference in TAD usage patterns is not simply an artifact of tumor cells displaying a higher number of expressed genes.

As a caveat, we note that our analysis can only infer correlation, not causation. While mutations leading to the mis-specification of TAD boundaries have been associated with certain cancers [15–17], there are diverse epigenetic mechanisms underpinning tumorigenesis [40–42] and the increased expression clustering we observe in the TADs of tumor cells could either be a cause or an effect of these mechanisms.

We next sought to understand changes in TAD usage during cell differentiation, a process where two opposing gene expression trends are in play: an overall decline in the number of genes expressed [19], but also an increase in the expression of specific lineage-determining genes [43]. The scRNA-seq transcriptomes described above are from single-cell assays covering cell differentiation in a variety of tissues. Limiting ourselves to assays with a single, well-defined differentiation trajectory, we found that in most assays (21 of 24, **Figure 4D**), cells earlier in the differentiation process exhibited greater expression clustering into TADs (*p* = 0.00019, one-sided t-test). These results are robust to how early vs. late differentiation is demarcated (**Methods**, **Figure S3**). Notably, our finding that both undifferentiated cells and tumor cells show greater expression clustering into TADs mirrors the well-documented transcriptional similarities between stem cells and tumor cells [44].

We then tried to resolve which of the two processes — broad-based gene deactivation or lineage-specific gene activation — was primarily responsible for the expression-clustering differences between early and late stage cells. For the scRNA-seq datasets described above, we compared the relative prevalence of low and high usage TADS across the stages, finding the most substantial change to be an increase in low-usage TADs (specifically, those with just one expressed gene) in late stage cells (**Figure 4F**, **Methods**). We interpret this as evidence for activation of new genes in hitherto-unused TADs as being the dominant factor behind the change in clustering patterns of expressed genes: if deactivation of previously active genes had instead been the dominating factor, fewer high-usage TADs in late stages should have been observed (since high-usage TADs contain more active genes). In contrast, activation of a specific lineage-determining gene in a hitherto inactive TAD would lead to a greater number of such single-usage TADs in late-stage cells, explaining our observations. Previously, the activation of lineage-determining genes in hitherto inactive TADs has been reported in mouse neural differentiation [21]). Our findings suggest such activation may be a broad phenomenon in cell differentiation.

Analysis of RNA velocity data provided additional support for this interpretation. In these datasets, both unspliced and spliced mRNA transcripts of a gene are available and comparing their relative abundance allows us to infer the recency of the gene’s transcription (“RNA velocity”), thus reliably reconstructing if a gene is being activated or deactivated [45]. Applying the RNA velocity tool *scVelo* [46] on two single-cell datasets (endocrine development in pancreas [47] and dentate gyrus development [48]), we estimated genes being activated and deactivated in each cell (**Figure 4G,H**). We then applied a bootstrap test (similar to the one described previously) to investigate if these genes were under- or over-dispersed across TADs (**Methods**), finding that in the case of gene activation, the set of genes was spread over significantly more TADs than would be expected by chance (mean z-score = 3.97, *p* ≈ 0, *t* statistic = 258, one-sample t-test). In contrast, the evidence for over-dispersion of deactivating genes was less strong, especially in the dentate gyrus data; however, it was still significant (mean z-score = 1.22, *p* ≈ 0, *t* statistic = 90, one-sample t-test).

### TAD Signatures: interpreting single-cell RNA-seq data with the TAD Map

The unprece- dented detail and volume of data from single-cell RNA-seq (scRNA-seq) assays has had a transformative impact on research in cellular, developmental, and translational biology. However, due to data sparsity, uneven gene coverage (“dropouts”), and protocol-specific variations in such datasets [49], the noise and error in scRNA-seq observations remain an impediment in drawing more accurate biological inferences. We introduce *TAD signatures*, a TAD-based representation of scRNA-seq data that can be used to augment standard scRNA-seq analyses with more robust input representations. We define a cell’s TAD signature as a feature vector of length *n* (= number of TADs in the scaffold that contain at least one gene, i.e., capacity ≥ 1), with the *i*-th feature quantifying the expression activity of TAD *i* on a scale from 0 (inactive) to 1 (fully active). We emphasize that TAD signatures can be inferred from just scRNA-seq counts by using the species-level TAD Map— no Hi-C data is needed.

To infer TAD signatures from a single-cell dataset, we designed a probabilistic generative model of TAD activation, modeling the activation state of each TAD in a cell as a Bernoulli random variable (0=“OFF”, 1=“ON”). The observed expression of a gene in a TAD is then modeled as a Poisson random variable, with the rate parameter (*λ*) depending only on the encompassing TAD’s state. Our model allows for a low rate of expression (*λ_OFF_*) even in TADs designated as “inactive” in order to robustly handle cases where a single gene might be active in an otherwise inactive TAD and to accommodate the noise and sparsity in single-cell datasets (**Methods**). Since the coverage depth and gene counts of scRNA-seq datasets vary, the observed rate parameters (*λ_ON_, A_OFF_*) will be specific to each dataset. Accordingly, we applied an expectation maximization (EM) algorithm to fit the parameters of a two-component mixture model to a scRNA-seq dataset, simultaneously estimating the two Poisson rate parameters and the activation probabilities of TADs in each cell. These activation probabilities comprise the TAD signature. For a single-cell dataset with *m* cells, the TAD signature thus consists of *m* feature vectors.

We believe that using TAD signatures to augment raw transcript count information is a broadly applicable approach for enhancing the biological signal in any scRNA-seq dataset. One interpretation of TAD signatures is as an epigenetically-informed dimensionality reduction of the scRNA-seq data. Alternatively, they may be viewed as infusing prior knowledge of chromatin structure into scRNA-seq analysis. They apply our finding that expressed genes are often clustered into TADs, robustifying expression estimates by averaging over gene groups within which higher levels of coexpression are expected *a priori* (by virtue of the shared TAD membership of genes in each group).

To assess the usefulness of TAD signatures in downstream scRNA-seq analyses, we first evaluated them on cell type inference, an important task in scRNA-seq analysis that continues to require extensive manual oversight. Applying the Leiden algorithm for cell clustering, we compared the biological accuracy of the clustering results computed from three input representations of scRNA-seq observations: raw RNA-seq data (generated by applying principal component analysis, PCA, on transcript count data); TAD signatures (by applying PCA on log-odds of TAD activation probabilities); and a concatenation of the two. On three large scRNA-seq datasets sourced from the Chan-Zuckerberg Biohub’s cellXgene portal [50], covering breast [51], lung [52] and T cells [53], we compared the overlap between the automatically-computed cell clusters and the expertly-annotated cell type labels made available by the original study authors (**Figure 5B**, **Methods**). We found that using TAD signatures led to cell clusters that better agreed with expert labels and in all cases, the representation that combined raw RNA-seq and TAD signatures performed at least as well as the RNA-seq-only representation.

**Figure 5:**
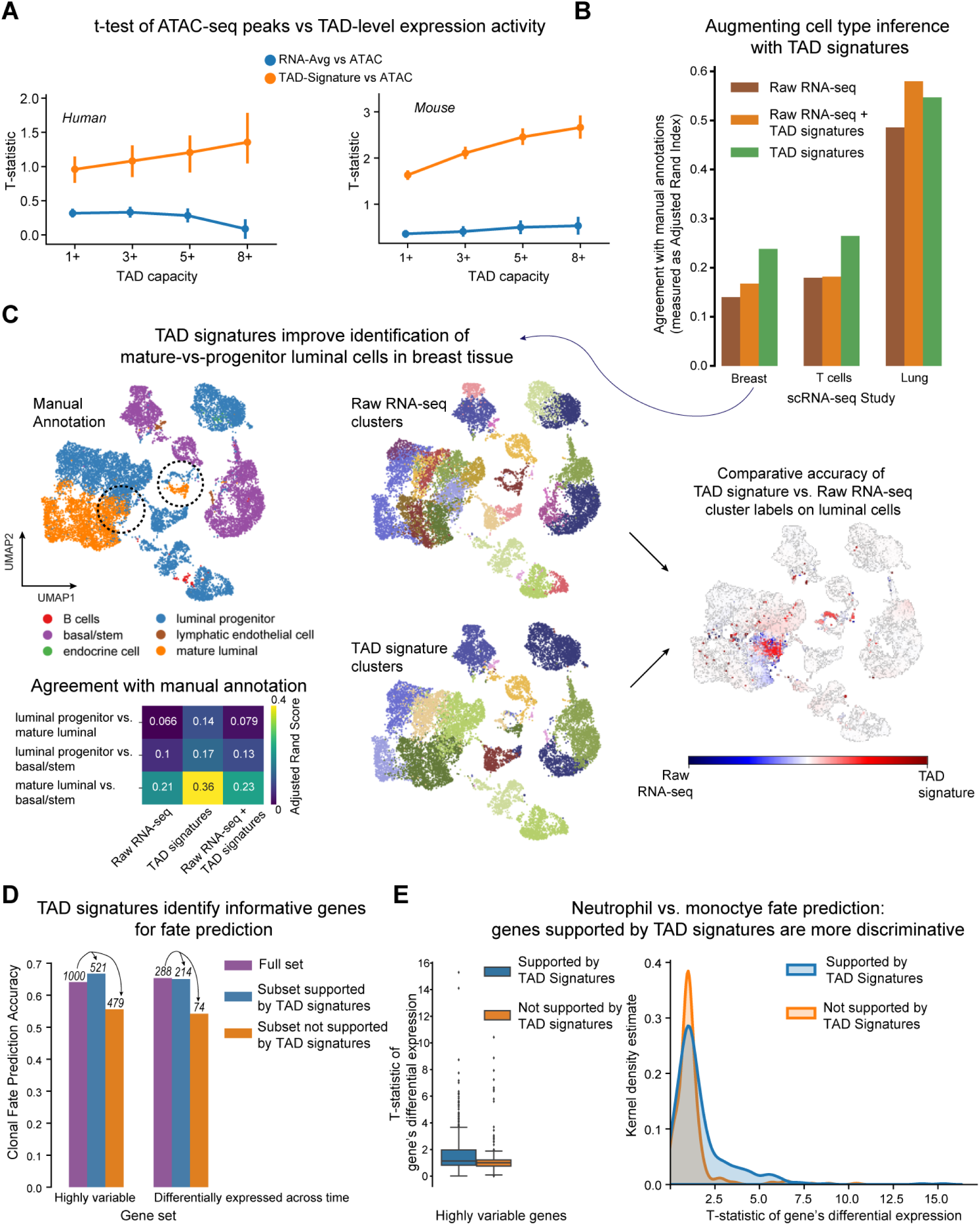
**TAD signatures augment standard scRNA-seq analysis**: We define a cell’s TAD signature, computed by overlaying its RNA-seq data on the TAD Map, as the set of per-TAD scores indicating the probability that genes in the corresponding TAD are transcriptionally active. **A**) On two multimodal datasets (paired scRNA-seq & scATAC-seq), the TAD signature, which summarizes expression activity in a TAD, shows greater agreement with ATAC-seq modality than simply averaging the gene expression in each TAD. The y-axis shows the t-statistic of average gene expression (“RNA-Avg”) of genes in a TAD or the TAD’s signature score (“TAD-Signature”), conditioned on observing any ATAC-seq peaks within the TAD’s boundaries (**Figure S4** shows an alternate evaluation). T-statistics were computed independently for each TAD and error bars indicate the 95th percentile range of this distribution. **B**) TAD signatures facilitate greater accuracy in automated cell-type inference: compared to Leiden clusters computed using just scRNA-seq data, Leiden clusters computed using only TAD signatures or from a combination of scRNA-seq and TAD signatures results in a closer match to manual annotations of cell types on three different scRNA-seq studies. For breast and T cell data, using the TAD signature alone is better than combining it with RNA-seq, possibly due to the latter being especially noisy in these datasets. Relatedly, we note the lower overall ARI for these datasets. **C**) A deeper study of the breast tissue data suggests that TAD signatures enable more accurate distinction between progenitor and mature luminal cells, indicating the usefulness of TAD signatures in studying cell differentiation. The colors in the middle-column plots correspond to clusters inferred using Leiden clustering; compared to TAD signature-based clusters, RNA-seq resulted in more clusters and thus colors. **D**) On data from Weinreb et al.’s lineage tracing study of hematopoiesis that also assayed scRNA-seq for all cells [56], we sought to predict the dominant fate of a clonal lineage (at days 4 or 6) using transcriptomes of same-clone sister cells harvested on day 2. In the original study, the best performance was achieved by using a set of highly variable genes or those differentially expressed across time. Using TAD signatures enables us to identify the more informative subsets of these genes, done by selecting genes only from highly variable TADs. **E**) We then quantified the discriminative power of each gene for distinguishing between the two most common cell fates (neutrophils and monocytes), measuring the gene’s t-statistic of differential expression between the two fates, and found that genes with support from TAD signatures have a higher discriminative power.

Seeking to understand why TAD signatures provided additional discriminative power, we drilled deeper into the breast tissue data. A particular challenge there is distinguishing between progenitor and mature luminal cells: there are some sub-populations of cells that are transcriptionally similar overall, but where Bhat-Nakshatri et al.’s expert knowledge of relevant marker genes allowed them to distinguish between progenitor and mature luminal cells [51]. We found that TAD signatures were particularly helpful in these ambiguous situations, enabling more accurate distinction between mature luminal cells and progenitor cells (**Methods**, **Figure 5C**). Such cases are often problematic for automated cell type inference methods since the distinction between two cell types may rest on subtle transcriptional differences that are masked by an overall similarity between their transcriptional profiles. Since TAD signatures are based on gene groupings informed by chromatin structure, they may help accentuate the epigenetic changes that accompany differentiation, allowing a clearer signal than would be available just from raw RNA-seq data. Thus, the incorporation of TAD signatures into scRNA-seq analysis pipelines may improve cell type identification.

Recently, it has become possible to simultaneously acquire multiple modalities at a single-cell resolution. While these technological advances promise novel insights, they also present new analysis challenges as these modalities— whose data is often sparse and noisy— need to be synthesized. Focusing on Cao et al.’s sci-CAR data from paired RNA-seq and ATAC-seq assays [54], we evaluated if TAD signatures can aid the analysis of such multimodal data. There, relatively low correlations have been observed between per-cell RNA- and ATAC-seq observations [55], despite chromatin accessibility around a gene being a fundamental determinant of its expression. This is likely due to the sparsity of ATAC-seq data (each peak can be present only 0, 1, or 2 times per cell), leading to noisy estimates [55]. We reasoned that TAD signatures will aggregate RNA-seq data in genomic intervals that are relevant to ATAC-seq data, making the agreement between the modalities clearer. Towards this, we aggregated ATAC-seq peak counts in each TAD and compared them against two measures of TAD-level gene expression: a) the TAD signature, and b) average RNA-seq expression of genes in the TAD. We applied two tests to quantify this agreement (**Methods**): a t-test measure that treats peak presence as a binary event (**Figure 5A**) and a Spearman rank correlation of peak counts with TAD activation probabilities or average RNA-seq counts (**Figure S4**). In both cases, we found that TAD signatures agreed substantially more with ATAC-seq peak data than the raw RNA-seq averages, suggesting that TAD signatures may also be useful in multimodal analyses.

Having documented changes in TAD usage during cell differentiation, we wondered if TAD signatures could inform cell fate prediction, i.e., the task of predicting the terminal type of a partially-differentiated cell. We applied them to Weinreb et al.’s single-cell lineage-tracing RNA-seq study of hematopoiesis [56]. In it, each progenitor cell (at day 0) was marked with a lentiviral construct that delivered an expressed heritable barcode, allowing clonal lineages to be traced across differentiation stages. To measure the power of RNA-seq data for cell fate prediction, the original study predicted the dominant cell type of a clonal lineage (inferred from fully-differentiated cells harvested at days 4 and 6) from the scRNA-seq transcriptomes of partially-differentiated cells of the same lineage (cells harvested on day 2). Weinreb et al. had found that the most informative genes for this prediction were differentially expressed or highly variable genes.

Using TAD signatures, we were able to further hone in on informative genes, achieving prediction accuracy comparable to the original study but with substantially fewer genes (**Figure 5D,E**). We did so by simply excluding genes that occur in TADs whose activation remains stable across the differentiation process, i.e. the TAD’s activation probability is not highly variable (**Methods**). This would happen if the gene is in a high capacity TAD where the activation state of other genes remains mostly unchanged throughout the dataset. Intuitively, this can be interpreted as an anomaly filtering step that removes genes whose observed transcriptional variability is inconsistent with other genes that co-occupy the TAD. On repeating the original study’s logistic regression analysis after this filtering, we were able to reduce the highly variable gene set by 47.9% (479 of top 1000 variable genes) and the differentially expressed gene set by 25.7% (74 of 288) while achieving similar or higher prediction accuracy (**Figure 5D**).

To further confirm that TAD signatures can indeed help identify informative genes, we quantified each gene’s informativeness for distinguishing between neutrophils and monocytes, the two most common terminal cell types in the study. Limiting ourselves to clonal lineages with these terminal cell types, we computed a t-statistic to quantify the separation between the day 2 (i.e., partially differentiated cells) expression values of the gene in neutrophil and monocyte lineages, finding that genes supported by TAD signatures had higher discriminative power than the rest (median t-statistic of 1.17 vs. 1.0; 75*^th^*percentile t-statistic of 1.97 vs. 1.22, **Figure 5E**).

### TADs may provide an environment of shared transcriptional control

Seeking to understand why gene expression clusters into TADs, we wondered if TADs facilitate co-transcription of neighboring genes. Specifically, we hypothesized that in a transcriptionally active TAD the chromatin structure enables transcriptional machinery to easily “move over” to the next gene, i.e., stochastically co-transcribe adjacent genes; we note that this hypothesis does not preclude other mechanisms by which chromatin folding regulates transcription [11]. As we have already documented, adjacent gene pairs which share a TAD are co-expressed more strongly than pairs that do not (**Figure 3F**). Seeking additional evidence, we compared adjacent gene pairs that have the same orientation (i.e., their transcription start sites (TSS) are on the same strand) to adjacent gene pairs that have opposite orientations, reasoning that if adjacent gene pairs are being stochastically co-transcribed, it should be easier for the transcriptional machinery to “move over” to the next gene if its TSS is correctly oriented and accessible (i.e., the genes have the same orientation). Remarkably, we found this to indeed be the case: analyzing RNA-seq data, the frequency of intergenic transcripts, and synteny between human and mouse, we found similarly-oriented adjacent genes in TADs to be significantly more co-expressed and evolutionarily preserved than opposingly-oriented pairs; this dependence on orientation is not as strong when evaluated on adjacent gene pairs that do not share a TAD.

Analyzing bulk RNA-seq datasets (same as in **Figure 3F**), we found that within TADs, gene pairs with the same orientation demonstrated significantly higher co-expression than opposingly-oriented gene pairs (Pearson correlation of 0.19 vs 0.15 for human and 0.22 vs 0.17 for mouse, with *p <* 10^−11^ in both cases, one-sided t-test). In contrast, for adjacent gene pairs that do not share a TAD, their relative orientation did not make a significant difference to the level of their co-expression (**Figure 6A**).

**Figure 6:**
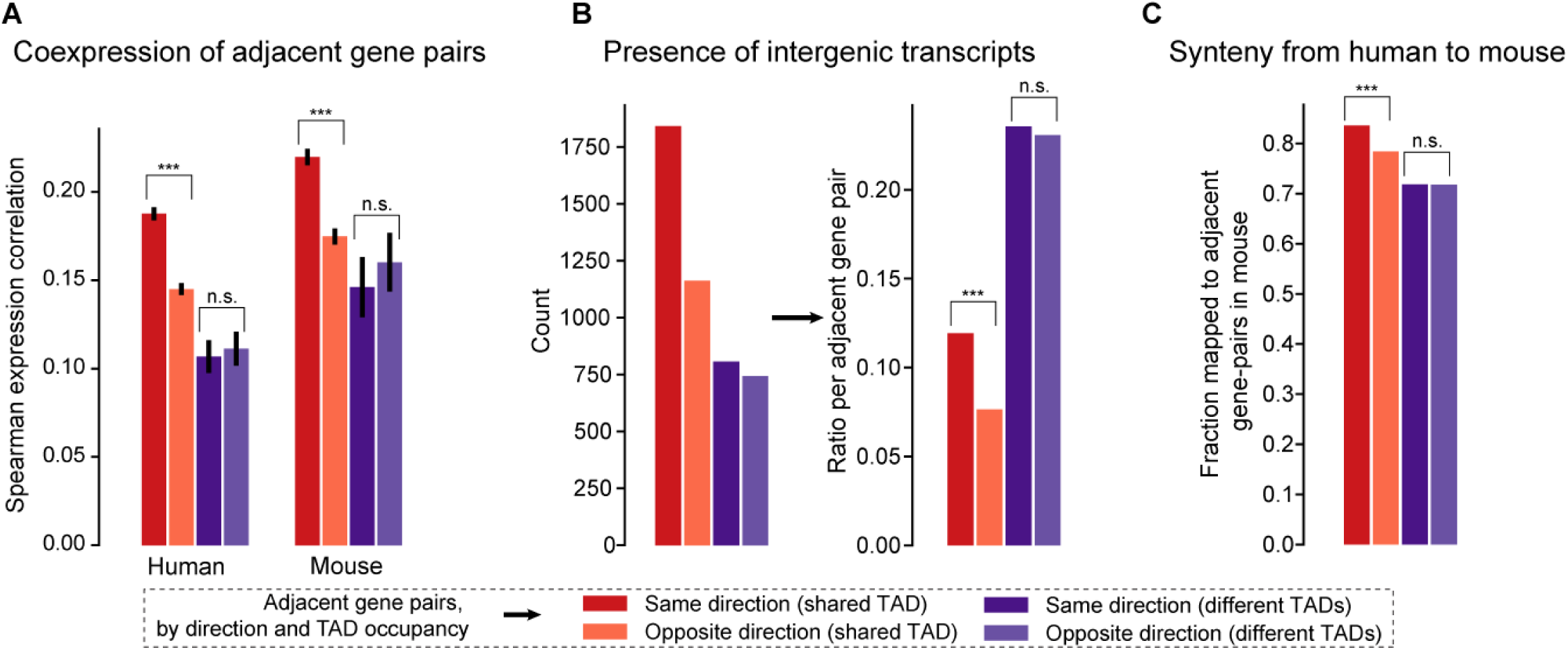
**Importance of gene orientation inside TADs suggests shared transcriptional control**: **A**) Extending the analysis from Figure 3F, coexpression between pairs of genes located adjacently on the genome is shown, now further split by whether the two genes have the same or different orientations (i.e. divergent or convergent). For gene pairs that share a TAD, having the same orientation is associated with significantly higher coexpression (*p <* 0.001, one-sided t-test); this is not the case for genes that do not share a TAD. **B**) Frequency of likely-erroneous intergenic transcripts (these exclude non-coding RNAs) between adjacent gene pairs is shown. Within TADs, intergenic transcripts are significantly more likely between same-direction gene pairs (*p <* 0.001, one-sided t-test), consistent with their co-transcription. Overall, however, the frequency of intergenic transcripts is higher outside TADs, possibly due to higher transcriptional error rates there. **C**) Synteny of adjacent gene pairs from human to mouse: adjacent gene pairs that shared a TAD in human are significantly more likely to have orthologs *adjacently* located in mouse. Additionally, gene pairs oriented in the same direction in a TAD are even more likely to be conserved syntenically (*p <* 0.001, one-sided t-test).

We also reasoned that if TADs provide an environment where the transcriptional machinery can “move over” to adjacent genes, the frequency of erroneous intergenic transcripts between adjacent genes should reflect this (these transcriptional mistakes are later degraded by the error-checking cellular machinery like the protein XRN2 or the exosome [57]). Analyzing a dataset of intergenic transcripts curated by Agostini et al. [57], we characterized each adjacent gene pair by the number (and proportion) of distinct intergenic transcripts between the genes (**Methods**, **Figure 6B**). We found that within TADs, similarly-oriented gene pairs had significantly more intergenic transcripts than adjacent pairs with opposing orientations (per-pair frequency of 11.9% vs. 7.6%, *p* = 1.03 × 10^−77^, one-sided binomial test). In contrast, gene orientation did not impact the frequency of intergenic transcripts between adjacent gene pairs that do not share a TAD. Notably, we did observe an overall greater prevalence of intergenic transcripts in latter gene pairs, suggesting a higher overall error-rate in transcription in non-TAD regions.

Lastly, we evaluated the extent to which these gene pairs are syntenically conserved from human to mouse. Sourcing orthology information from Ensembl, we queried adjacent gene pairs in human that had mouse orthologs, checking if the orthologs were also adjacently located in the mouse genome (**Figure 6C**). Consistent with the functional importance of TADs, we found within-TAD gene pairs to be more syntenically conserved than gene pairs that do not share a TAD (81% vs. 71.8%, *p* = 6.6^−40^, one-sided binomial test). Furthermore, within TADs, pairs with the same orientation were preserved significantly more than those with opposing orientations (83.6% vs 78.4%, *p* = 1.6 × 10^−30^, one-sided binomial test). To summarize, there is thus evolutionary and expression-based evidence that co-transcription of adjacent genes in TADs is more likely if the adjacent gene pairs have the same orientation, suggesting that a stochastic carry-over of transcriptional machinery may be at work.

## Discussion

We have leveraged Hi-C data from a limited number of cell types to estimate a consensus TAD scaffold that we show to be informative in a species-wide context: across a variety of cell and tissue types, we document higher rates of eQTL prevalence inside our estimated TADs and higher coexpression of adjacent genes that co-occupy a TAD. From the TAD scaffold we then deduced the TAD Map, a representation of TADs as the set of genes contained in them. In designing this lower-resolution representation, we sought a balance between preserving enough detail for the abstraction to be informative while avoiding assumptions that may not hold across cell types. Accordingly, though we believe our inferred TAD boundaries to be broadly accurate — they are supported by CTCF-binding data in a variety of tissues — our transcriptional analysis relies only on interpreting the TAD Map as a grouping of genes along the genome, thus making it more robust to the precise locations of TAD boundaries.

The TAD Map’s gene grouping is based on the genome’s overall chromatin structure, enabling our investigation of the latter’s influence on transcription. We find that expressed genes in a cell are likely to cluster into TADs, with this expression clustering significantly stronger in undifferentiated and tumor cells. Our finding systematizes and is consistent with previous cell type-specific reports of TADs being a unit of epigenetic control of transcription [8–11]. As the cell differentiates, we found this expression clustering to get weaker, with newly expressed genes likely to be activated in a more TAD-independent manner. Bonev et al. [21] observed that neural differentiation in mouse was marked by hitherto inactive TADs becoming active; our analysis suggests such activation of lineage determining genes in previously inactive TADs may be a hallmark of cell differentiation. Our work may also inform studies on the regulatory range of transcription factors [43]. We note that a systematic approach that can cover a large corpus of RNA-seq data is crucial for the statistical power to discern subtle links between chromatin structure and transcription. Such patterns may not be evident otherwise: e.g., in an analysis limited to only two cell types, Long et al. [26] report not observing correlations of genes inside TADs beyond what gene proximity would suggest.

A key advance facilitated by the TAD Map is the ability to improve the modeling of scRNA-seq data by epigenetically-informed gene groupings. Any scRNA-seq dataset can be augmented with TAD signatures, our probabilistic representation that aggregates gene expression by TAD to estimate per-TAD activation probabilities. Intuitively, TAD signatures can be thought of as biologically-motivated dimensionality reduction of scRNA-seq data that summarize noisy, sparse transcript counts into fewer, more-informative features, analogous to principal component analysis or non-negative matrix factorization [49]. We have shown that they enable more accurate cell type inference, facilitate better agreement between paired RNA-seq and ATAC-seq data, and identify the genes most informative in predicting cell fate. More broadly, we believe a promising research direction for future work is to systematically leverage our growing insights into chromatin structure (e.g., its hierarchical organization) to build likelihood models of gene coexpression that can be integrated into the standard analysis pipelines of scRNA-seq data.

Characterization of the broad principles that link transcription to chromatin’s folding structure still remains an open question. While we have focused only on the overall structure of chromatin that would hold across cell types, minor variations in chromatin structure occur at a cellular level and those variations would result in transcriptional differences not captured by the TAD Map. Also, chromatin folds hierarchically, with compartments, TADs, and sub-TADs being successively smaller units of folding. We limited ourselves to TADS, the most well-studied tier, but future work could build towards more nuanced TAD Maps that are cognizant of this hierarchy. In our inferred TAD scaffold, genes can cross TAD boundaries, sometimes spanning two TADs. We associated such genes with both TADs, which is the conservative choice from a clustering test perspective; such genes could be further investigated. It may also be beneficial to jointly infer TAD Maps across species. Compared to human, fewer Hi-C datasets are available for mouse and though we sought to compensate for this with appropriate hyperparameter choices in our maximum likelihood estimation, our predicted TAD scaffold for mouse nonetheless includes some large TADs (containing over fifty protein-coding genes), bigger than those in human.

We see the TAD Map and its potential for “transcriptional imputation” in cell/tissue studies as akin to the role of genotype imputation for genome-wide association studies.

## Methods

### Cell type-specific TAD definitions

We sourced cell type-specific TAD architectures from Liu et al.’s TADKB database [31], selecting TAD definitions inferred with the Directionality Index (DI) technique at 50 Kb resolution. We chose TADKB because it ensured a consistent “Hi-C to TAD” mapping across experimental data from multiple studies. We retrieved data for all the human and mouse cell lines available in the database: seven for human (GM12878, HMEC, NHEK, IMR90, KBM7, K562, and HUVEC) and four for mouse (CH12-LX, ES, NPC, and CN). These TAD definitions were computed by Liu et al. using source Hi-C data from Rao et al. [7] (Gene Expression Omnibus, GEO, accession GSE63525) and Bonev et al. [21] (GEO accession GSE96107).

### Reference genome versions

All genomic coordinates and gene names correspond to Ensembl v102, with the human and mouse reference genomes being hg38 and mm10, respectively. With the *liftOver* program [58], TAD definitions for human cell types were mapped to the hg38 reference genome from hg19, the reference genome used in TADKB. TAD definitions for mouse cell types already corresponded to the mm10 reference genome.

### Maximum likelihood estimation of the consensus TAD scaffold

We infer TADs independently for each chromosome: our algorithm takes as input a list of cell type-specific TAD architectures for the chromosome and outputs the consensus TAD definition. Both the input and output definitions specify TADs as a set of non-overlapping genomic intervals along the chromosomes. Our algorithm currently operates at a 50 kb resolution with both the input and output defined at that granularity.

We divide the entire chromosome into segments of size R (currently, R = 50 kb) and for every possible pairwise combination of segments, compute the likelihood that the two segments share a TAD. More formally, given segments at loci *i* and *j*, we define *X_ij_* as the number of input cell type-specific TAD architectures in which *i* and *j* share a TAD. Our goal is to infer *C_ij_*, where *C_ij_* ∈ {0, 1} indicates if *i* and *j* share a consensus TAD. Additionally, *C_ij_* need to obey integrity constraints that correspond to a valid TAD architecture; we describe these later. To accommodate the differing amounts of input data available for human and mouse in a unified framework, we discretized *X_ij_* into three levels: *X_ij_* ≥ *T*, *T > X_ij_ >* 0, and *X_ij_* = 0. With data for 7 human cell types and 4 mouse cell types, we chose *T* as 4 (for human) and 2 (for mouse), so that the three levels of *X_ij_* express the intuition that it receives support from the majority, at least one, or none of the input cell types, respectively. We then parameterize our likelihood model *P*(*X_ij_* |*C_i j_*) as follows:

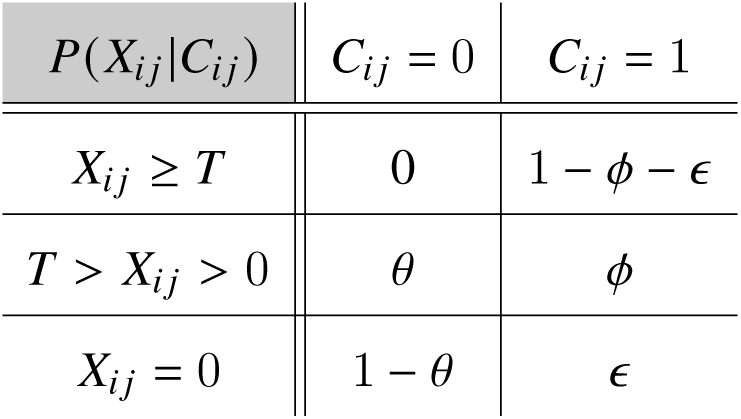

where 0 *< (), ¢, E,* (1 − *¢* − *E*) *<* 1. Our parameterization implies that a segment pairs *i, j* which share a TAD in a majority of the input cell types (*X_ij_* ≥ *T*) must be present in a consensus TAD. On the other hand, we provide for the possibility (*ε >* 0) that a segment pair *i, j* with no support from any of the input architectures might still share a consensus TAD— this allows us to stitch together overlapping TAD ranges across cell type-specific inputs if strong overall support exists for a broad TAD at that locus.

Under our model, the likelihood of the observations **X** = {*X_ij_* } given the consensus TAD scaffold **C** = {*C_ij_* } is

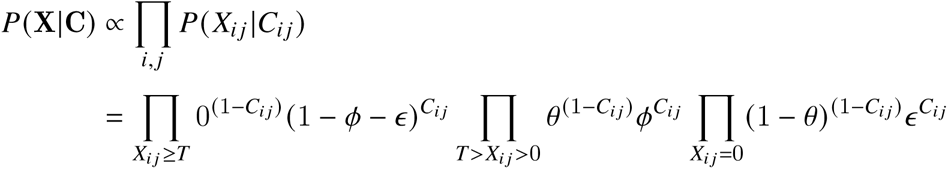

The first sub-product imposes the binding constraint that *C_ij_* = 1 for all segment pairs {*i, j* | *X_ij_* ≥ *T* }. Given that, we can focus on the remaining two sub-products to maximize the log-likelihood:

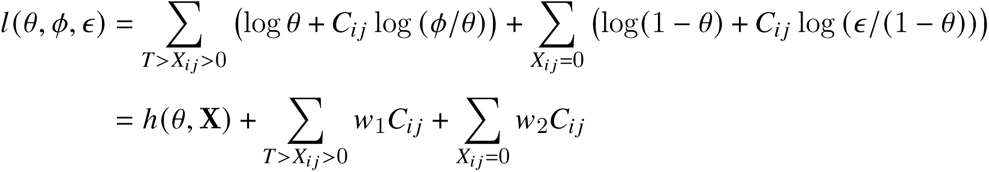

where *h*((), **X**) is not a function of *C_ij_*, and the terms *w*_1_and *w*_2_represent more convenient combinations of the parameters *(), ¢*, and *E* . Maximizing the log-likelihood thus requires solving

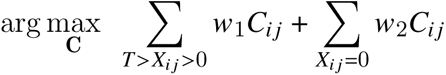

We note that integrity constraints on *C_ij_* link the terms: **C** needs to be transitive, i.e., if segments pairs (*i, j*) and (*j, k*) share a TAD then so must (*i, k*). Also, a TAD must be contiguous: if *C_ij_* = 1 then *C_iv_* = *C_v j_* = 1 for all segments *v* between *i* and *j* . Intuitively, this formulation describes a trade-off between biasing towards long TADs, which cover more true positive *C_ij_* s (guided by *T > X_ij_ >* 0 cases) but also have more false positives (driven by *X_ij_* = 0 cases), and short TADs where the false positives will be fewer but the risk of false negatives increases.

To maximize *1* and compute **C**, we formulated a dynamic programming algorithm that splits the chromosome into recursively smaller ranges and finds the globally optimal combination of *C_ij_* assignments. We chose *w*_1_= 0.5*, w*_2_= −1 for human and *w*_1_= 0.05*, w*_2_= −3 for mouse; these choices produced consensus TAD scaffolds where the number of TADs and the distribution of their lengths was in line with the corresponding statistics for cell type-specific TAD architectures.

### Expression quantitative trait loci

From the GTEx Analysis Release V8 [32], we sourced data for all 49 tissues with available single-tissue eQTL data. We filtered the data, limiting ourselves to eQTLs with p-value less than 10^−5^. For each gene and eQTL pair, we computed the genomic distance between the transcription start site (TSS) of the gene and the eQTL locus; based on it, we assigned the (TSS, eQTL) pair to one of the genomic-distance bins partitioned by the following cut-points (all units in kb): [0, 5, 10, 20, 30, 50, 75, 100, 150, 250, 350, 450, 550, 650, 750, 1000].

We counted the number of observed pairs in each bin, separately tracking pairs where the TSS and eQTL loci shared a TAD and pairs where they did not; we additionally separated pairs where the eQTL locus was upstream of TSS from those where it was downstream. The probability *p* (*d*) of an eQTL occurring at a distance *d* from the TSS was then estimated by dividing the number of observed (TSS, eQTL) pairs in each bin by the midpoint of the bin’s genomic range.

### Agreement with CTCF ChIP-seq

We sourced CTCF ChIP-seq data from ENCODE, acquiring all the available ChIP-Seq assays for human and mouse. We filtered to only keep peaks with Irreproducible Discovery Rate (IDR) less than 0.05 and mapped these peaks to our estimated TAD scaffold. We then grouped the peaks by their location relative to TADs, partitioning the entire genome into the following disjoint categories: *TAD Boundary* (50 kb segments on each side of the TAD), *Inside Boundary* (two 50 kb segments inside the TAD, just interior to the TAD boundaries), *Outside Boundary* (two 50 kb regions outside the TAD, just exterior to the TAD boundaries), *TAD Interior* (the part of the TAD that’s not in the *Inside Boundary* segments), and *TAD Exterior* (all other parts of the genome). With the CTCF peak widths being much smaller (median width = 273 bp) than the granularity of our TAD scaffold (50 kb resolution), we assumed that a peak would not span two segments and assigned each peak to its genomic segment based only on the peak’s midpoint locus. Finally, we counted the number of peaks in each segment category, normalizing that count by the aggregate length of genomic segments in that category.

### Bootstrap test for clustering of expressed genes into TADs

The bootstrap test operates on data from a single cell (in scRNA-seq data) or a single tissue (in bulk RNA-seq data). Given scRNA-seq readout from any cell, we compute *k*, the number of genes with non-zero transcript counts in the cell. Treating gene activity as a binary event, we then generate 500 bootstrap samples of single-cell gene expression in each of which we randomly choose *k* protein-coding genes to be active. For both the actual observation and the bootstrap samples, we map these genes to the TAD Map, computing the number of TADs *n*(*p, k*) which have *p* or more active genes; here, *p* = 1 corresponds to identifying the set of TADs with non-zero usage. We estimate the mean and standard deviation of the distribution *n*(*p, k*) from the bootstrap samples and, using that, compute the z-score for the actual observation. In bulk RNA-seq data, which includes gene expressed aggregate from a collection of cells, almost all genes have some non-zero expression. There, we pre-set a threshold *k* (say, 5000) and limit ourselves to the top-*k* genes by transcript count; the bootstrap test and z-score are computed for this *k*. **Figure S2B** shows that our results for bulk RNA-seq are robust to a range of choices for *k*.

In the test above, if a gene spans two TADs we count it in both TADs. This avoids us having to assign the gene to one TAD or the other arbitrarily. We also believe this to be the conservative choice for our clustering test: it will lead to more TADs per gene— i.e., lower clustering of expressed genes into TADs— than if we were to assign the gene to just one TAD or the other.

### scRNA-seq cell differentiation datasets

We sourced scRNA-seq data from the CytoTrace database made available by Gulati et al. [19]. The study collected and curated scRNA-seq cell differentiation studies across multiple species, covering a variety of protocols and tissues. We chose this corpus as our scRNA-seq testbed, since it covers a diversity of protocols and tissues and allowed us to extend our analysis of also study TAD usage during cell differentiation. Of the 43 datasets available on cytotrace.stanford.edu (while their webpage lists 47 entries, 4 rows are blank), we filtered out studies with fewer than 200 cells and those that did not originate from human or mouse tissue, leaving us with 33 studies. We had difficulty converting 3 of these from the original *RDS* format to a *Scanpy*-compatible format and limited ourselves to the remaining 30 (which covered 70,243 cells); these formed our scRNA-seq corpus.

### Designation of early vs. late-stage cells during differentiation

When analyzing TAD usage during cell differentiation, we further limited ourselves to the 24 scRNA-seq datasets where the putative differentiation trajectory did not have any branches, allowing us to reliably order cells along a differentiation time course. We used Gulati et al.’s annotations of differentiation stage in each study and considered two measures of early versus late differentiation: 1) consider only the cells at the first differentiation stage (*order* = 0 in CytoTrace) as “early”, with all other cells comprising the “late” stage, or 2) partition the cells in each dataset by *order* so that cells are divided roughly equally between the first (“early”) and second (“late”) halves.

### Comparison on TAD usage in early vs. late-stage cells

For the scRNA-seq datasets described above, we compared the relative prevalence of low (usage = 1) and high (≥ 3) usage TADS, seeking to ascertain whether lower levels of expression clustering in late-stage cells were because of a relative abundance of low-usage TADs or if they were due to a paucity of high-usage TADs. We found stronger evidence for the former than the latter, observing that both phenomena occur to a certain extent. Across each of the 24 datasets, we compared TAD usage between early and late stage cells (**Figure 4F**), finding a stronger signal in low-usage TADs: early stage cells had fewer low-usage TADs (mean early-vs.-late z-score difference = -0.36, *p* = 0.048, one-sided paired t-test). In contrast, the stages were less clearly separated in their z-scores corresponding to high-usage TADs (mean z-score difference between stages = -0.10).

### Analysis of RNA velocity data

We sourced the preprocessed *Scanpy*-compatible versions of the two datasets [48] from the RNA velocity tool *scVelo* (version 0.2.2) [46]. Applying *scVelo*, we computed 1) a pseudotime estimate for each dataset, allowing us to order cells along a time course, and 2) a velocity estimate for each gene in each cell. As activating genes we selected gene–cell combinations where the gene’s velocity in the cell was in the top (i.e., most positive) 2.5 percentile across all such combinations. To limit ourselves to cells where broad-based activation was in play, we removed from consideration cells with less than 15 genes marked as being activated. Deactivating genes were estimated similarly, with gene–cell combinations chosen if their velocity was in the bottom (i.e., most negative) 2.5 percentile across the overall data set.

### Intergenic transcripts

We sourced data from Agostini et al.’s study of intergenic transcripts. They collected and analyzed data from 38 publicly available datasets, covering over 2.5 billion uniquely mapped reads. We limited ourselves to 11,417 intergenic transcripts that are currently unannotated (the others corresponded most frequently to non-coding RNA fragments). As in the original study, each transcript was mapped to its adjoining genes as per the Gencode 27 reference. We further annotated each intergenic transcript by whether its adjoining genes were a) shared a TAD, and b) if they were oriented similarly.

### TAD signatures: probabilistic model

We define a 2-component mixture model to infer TAD activation probabilities of a single-cell dataset. Let *X* ∈ R*^n^* be the gene expression values for a single-cell dataset with *n* cells and *p* genes, with a particular cell *c*’s expression being *x*^(^*^c^*^)^∈ R*^p^*. We assume that transcriptionally active TADs (“ON”) correspond to a higher rate of per-gene expression while inactive TADs (”OFF”) correspond to lower rates of gene expression. As mentioned earlier, our model allows inactive TADs to also generate non-zero gene expression, albeit at a lower rate than the “active” TADs. Doing so increases our robustness to noise and allows for one-off gene expression in a TAD. In the mixture model, the probability of *x*^(^*^c^*^)^’s expression in *c* is

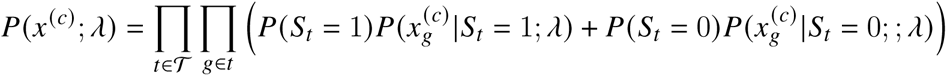

where *λ* = {*λ_ON_, A_OFF_*}; T is the set of all TADs with one or more genes; each TAD *t* ∈ T is a set of genes, with *g* being one such gene; *S_t_*is the Bernoulli random variable indicating the activation state of *t*, with *S_t_*= 1 or 0 corresponding to *t* being “ON” or “OFF”, respectively; and *x*^(^*^c^*^)^is the expression of gene *g* in the cell *c*. Here, T *, t* and the gene memberships in TADs are sourced from the species-specific TAD Map. We model that gene expression values are Poisson-distributed counts, though this assumption can be relaxed:

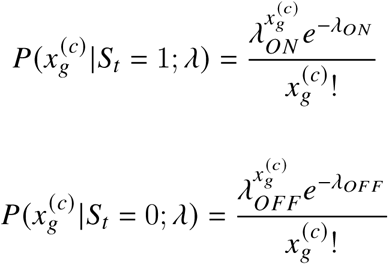

To infer the TAD signature for a particular dataset, we fit this model with the expectation maximization (EM) algorithm, seeking to maximize the log-likelihood over all cells:

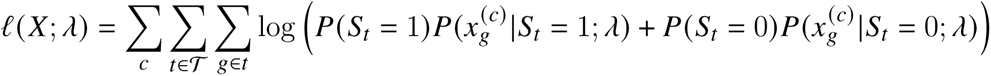

Since the maximum likelihood estimator of the Poisson rate parameter is just the sample mean, the implementation of the EM algorithm is simplified: each maximization round assigns *λ_ON_*and *λ_OFF_*as averages of observed gene expression values across all genes, weighted by the containing TAD’s activation probability *P*(*t* = 1). Also, we note that quasi-Poisson generalizations result in the same maximum likelihood estimator for the expected value of the rate parameter, suggesting that even in cases where the Poisson assumptions do not hold, the corresponding estimate is reasonable.

### Analyzing multimodal singe-cell data with TAD signatures

We retrieved single-cell multi-modal data from the sci-CAR experiment (GEO accession GSE117089), obtaining simultaneously-assayed chromatin accessibility (ATAC-seq) and gene expression (RNA-seq) modalities per cell. The sci-CAR dataset contains two sub-studies, one on 3,260 human cell-derived A549 cells and the other on 11,296 mouse kidney cells. The analysis we describe next was applied to each sub-study independently. We removed cells that had fewer than 20 genes with non-zero RNA-seq activity and also removed genes with RNA-seq activity in fewer than 10 cells. We count normalized (to 10^6^) both the RNA-seq and ATAC-seq modalities, and log transformed the RNA-seq modality. Gene identifiers were converted to Ensemble v102, genes and ATAC-seq “peaks” were mapped to the human TAD Map inferred in this work, and TAD signatures were estimated. For this analysis, we considered only genes and peaks that overlapped with a TAD.

Our analysis had dual, related objectives: assess if TAD signatures are a biologically meaningful summarization of gene expression and, secondly, evaluate if TAD signatures can be used to robustly synthesize gene expression data with other single-cell modalities. Towards this, we summarized gene expression activity in a TAD in two ways: i) the score for that TAD in the signature, corresponding to its activation probability, and ii) directly compute the average of expression values for genes in the TAD. We compared the agreement of these estimates to the ATAC-seq modality with two separate tests, both of which showed the TAD signatures to agree better with the ATAC-seq modality that averages of gene expression.

In the first test (**Figure S4**), for each TAD *t*, we computed the Spearman rank correlation of the cumulative ATAC-seq peak values in the TAD with each of the two expression estimates. In both species, we found TAD signatures to have a stronger correlation with ATAC-seq observations than simple averages of gene expression, suggesting that TAD signatures are a valuable summary of gene expression and can be useful in multimodal synthesis. However, the correlations were low overall, likely due to the sparsity in the ATAC-seq modality — over 99.5% of ATAC-seq peaks in a cell are typically empty [55].

We designed a second test to work around the sparsity issues with ATAC-seq data. For each TAD *t*, we considered the presence of any non-empty peak in it as a binary event and, dividing cells into two groups based on this, compared the expression between these groups by a 2-sided t-test. The intuition here is that cells with ATAC-seq activity in the TAD should have different expression than cells where no such activity was documented. Considering ATAC-seq activity as a binary event helps address the sparsity issues with it: the t-test is designed to work with populations (here, groups of cells) of unequal sizes, and is hence more robust to having relatively few cells with non-zero ATAC-seq values in the TAD. Computing t-statistics for each TAD, we compared the two expression measures (**Figure** 5A), finding that the t-statistics generated using the TAD signatures were substantially stronger than those from averages of gene expression values. Furthermore, the difference was starker for TADs with larger capacity (i.e., those that contain more genes). We believe this may be because arithmetic averages are prone to distortions by a single outlier gene while TAD signatures are more robust to such outliers due to their underlying probabilistic model.

### TAD signatures for cell type inference

We acquired single-cell data from the https://cellxgene.cziscience.com/ portal, obtaining *AnnData*-formatted [59] datasets from single-cell RNA-seq studies of the human lung (10x sub-study; European Genome-Phenome Archive accession EGAS00001004344; 65,662 cells; [52]), T-cells (GEO accession GSE126030; 51,876 cells; [53]), and breast epithelial cells (GEO accession GSE164898; 31,696 cells; [51]). Cells with fewer than 20 active genes and genes active in less than 10 cells were removed. For the breast tissue data, the dataset annotations seemed to suggest samples were grouped in two broad batches and, to reduce batch effects, we limited ourselves to the larger batch (17,153 cells). The data was then count normalized (to 10^6^) and log transformed using *Scanpy*. Gene identifiers were converted to Ensemble v102, genes were mapped to the human TAD Map inferred in this work, and TAD signatures were estimated. For each dataset, we then generated the following representations: i) principal component analysis (PCA) with 50 components, ii) log-odds 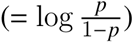 transformation of the TAD signatures computed on the dataset, followed by a 50-component PCA, and iii) a concatenation of the previous two representations. Leiden clustering [60] using *Scanpy* was performed on each of these representations. The datasets from cellxgene portal contained expert-annotated cell-type labels for each cell and we computed the adjusted Rand index (ARI) of the overlap between computed Leiden clusterings and the expert labels.

### Log-odds transformation of TAD signatures

We recommend a log-odds transformation when using TAD signatures to generate data representations for clustering, visualization, or predictive analysis. Many such analyses implicitly or explicitly rely on Euclidean distances between observations. The log-odds transformation converts probabilities (which are in [0, 1] range) to the full range of values in R, making it more amenable to such distance measures.

### Use of TAD signatures for cell fate prediction

We obtained data from Weinreb et al.’s lineage tracing study (GEO accession GSE140802; [56]), count normalizing and log transforming the transcript counts. No cell had fewer than 20 active genes and no gene was active in less than 10 cells. We limited our analysis to their *in vitro* observations, since the original study reported substantially greater predictability with this data than with their *in vivo* observations. We selected clonal lineages that had at least one cell harvested at day 2 (to use for making the prediction) and at least one cell harvested on day 4 or 6 that had been assigned to a differentiated cell-state (i.e., was not labeled “*Undifferentiated*”), resulting in 15,482 cells across 833 clonal lineages. Gene identifiers were converted to Ensemble v102, genes were mapped to the human TAD Map inferred in this work, and TAD signatures were estimated.

To predict cell fate from day-2 cells’ gene expression, we followed the original study in labeling each clonal lineage with the most frequently observed terminal cell type (day 4 or 6) of its cells. For a day-2 cell with gene expression *x* ∈ R*^n^* (*n* is the number of genes) and of clonal lineage C, a multi-class logistic regression sought to predict the cell-type label of C using *x*. *L*_2_-regularized logistic regression (using the Python package *scikit-learn* v0.24.1) was performed along the lines described in the original study, with observations corresponding to 1,209 day-2 cells split in training (49.9%) and test (50.1%) subsets. The test–train split was stratified by clonal lineage so that a clonal lineage was assigned only to the test or train subsets, but not both. The regularization hyperparameter (C=0.005) was chosen by a preliminary cross-validation analysis using the full set of genes. In all regressions below, the training and test sets as well as hyperparameters were kept unchanged.

For the task of lineage fate prediction from day-2 expression, Weinreb et al. had found greater predictive power by using only differentially expressed or highly variable genes for prediction. Following the original study, we used data on all cells (not just day-2 cells) to identify these gene sets. We constructed the set of differentially expressed genes by applying *Scanpy* (v1.4.6) to identify the top 100 most-differentially expressed genes corresponding to cells harvested on each of days 2, 4, and 6, respectively. The set union of these groups yielded 288 genes. Separately, the set of 1,000 highly variable genes was also computed with *Scanpy*, using default parameters. These gene sets formed the basis of our logistic regression analysis.

We quantified a specific TAD’s variability as *cr*^2^/*µ* where *µ, cr* are the mean and standard deviation of the TAD’s score (i.e., its activation probability) across the 15,482 per-cell TAD signatures; we note that the variance-to-mean ratio is a commonly used measure of dispersion for probability distributions. Sorting on this measure, we defined highly variable TADs as those in the top one-third. The above-described gene sets were each split into two, based on whether a gene was in one of these highly variable TADs. This resulted in a ‘214 (gene in highly variable TAD) + 74 (rest)’ split for the 288 differentially expressed genes and a ‘521 (gene in highly variable TAD) + 479 (rest)’ split for the 1,000 highly variable genes. The logistic regression analysis was repeated using each of these gene subsets (**Figure** 5C).

To assess the informativity of a specific gene, we limited ourselves to clonal lineages where the dominant terminal cell-type was either neutrophil or monocyte, allowing us to formulate a differential expression t-test. These are the two most common lineage fates in the data and the original study had also limited itself to comparing these lineages in some of their analyses. After the filtering, we grouped day-2 cells by their clonal lineage’s terminal cell type. For each gene, we applied a 2-sided Welch’s t-test to assess if it was differentially expressed between the two groups, finding that genes with high t-statistics were more likely to be located in highly variable TADs (**Figure** 5D).

### Quantification and Statistical Analysis

Statistical tests were conducted using version 1.3.1 of the SciPy Python package.

## Acknowledgements

We thank Anders Sejr Hansen, Brian Hie, and Stuti Khandavala for helpful discussions and technical assistance. RS and BB were partially supported by the NIH grants R01GM081871 and R35GM141861

## Author Contributions

All authors conceived of, contributed to, and wrote the paper. RS developed the method and implemented the software.

## Supplementary Materials

**Figure S1:**
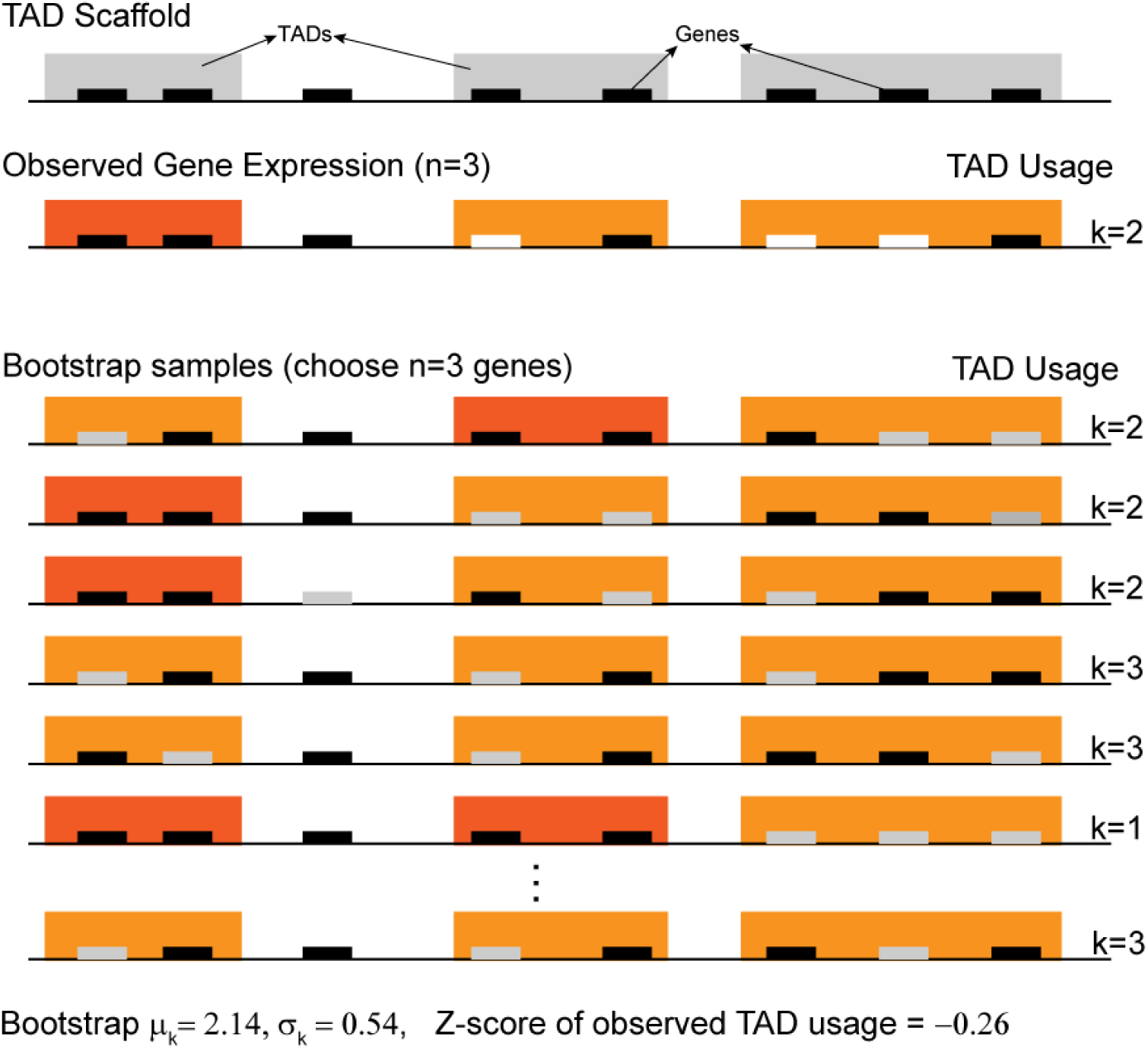
**Demonstration of bootstrap test for gene clustering**: This is an accompaniment to Fig 4A, showing the bootstrap test of gene clustering with a more complex, albeit still artificial, example.

**Figure S2:**
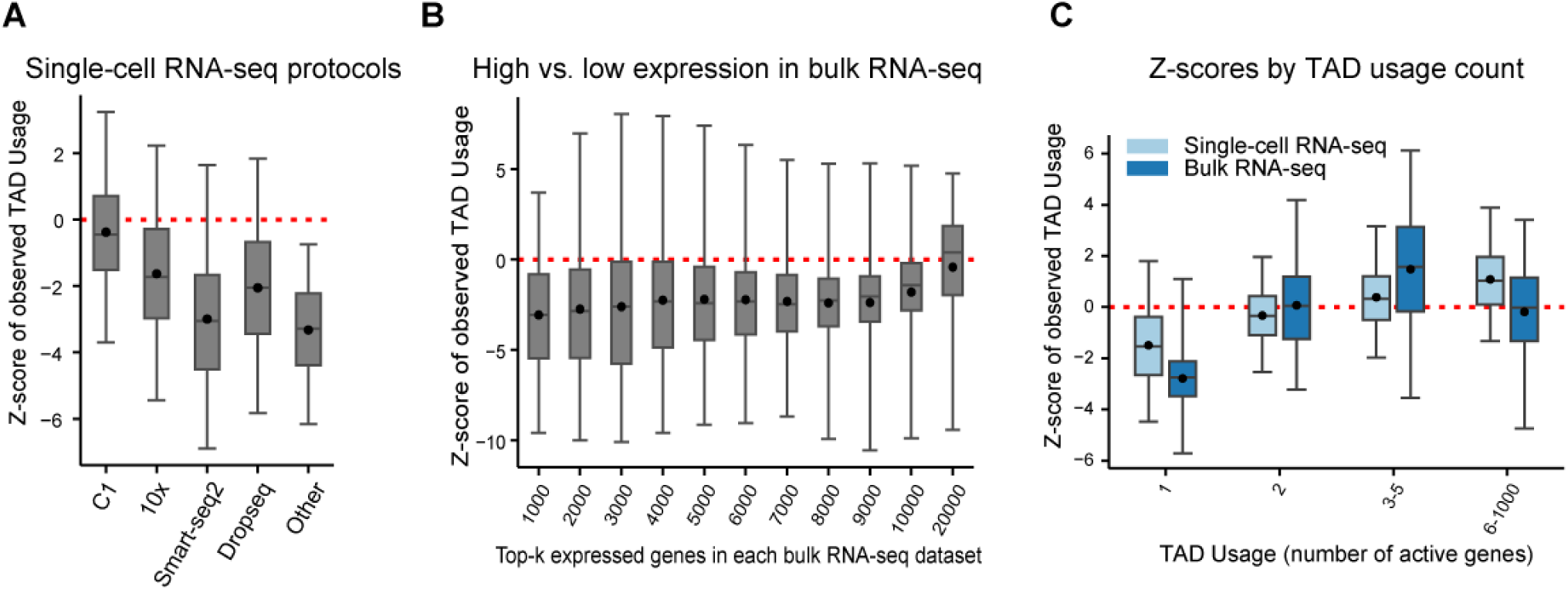
**Measurement of expression clustering into TADs is robust to various tests**: This figure accompanies Figure 4B-C where we analyzed 70,243 single-cell transcriptomes and 863 bulk RNA-seq datasets. **A**) This clustering is not an artifact of the single-cell RNA-seq protocol. **B**) In bulk RNA-seq datasets, where the greater library depth enables a deeper investigation, the clustering observation holds even for weakly expressed genes. We note that the last boxplot covers almost all protein-coding genes so that the bootstrap z-scores will be near-zero by design. **C**) The clustering effect is primarily due to an under-prevalence of TADs with just one active gene and partly due to an over-prevalence of TADs with 3 or more genes. In all boxplots in this figure, the box represents the 25-75*^th^*percentile range, and the whiskers represent 1-99*^th^*percentile range.

**Figure S3:**
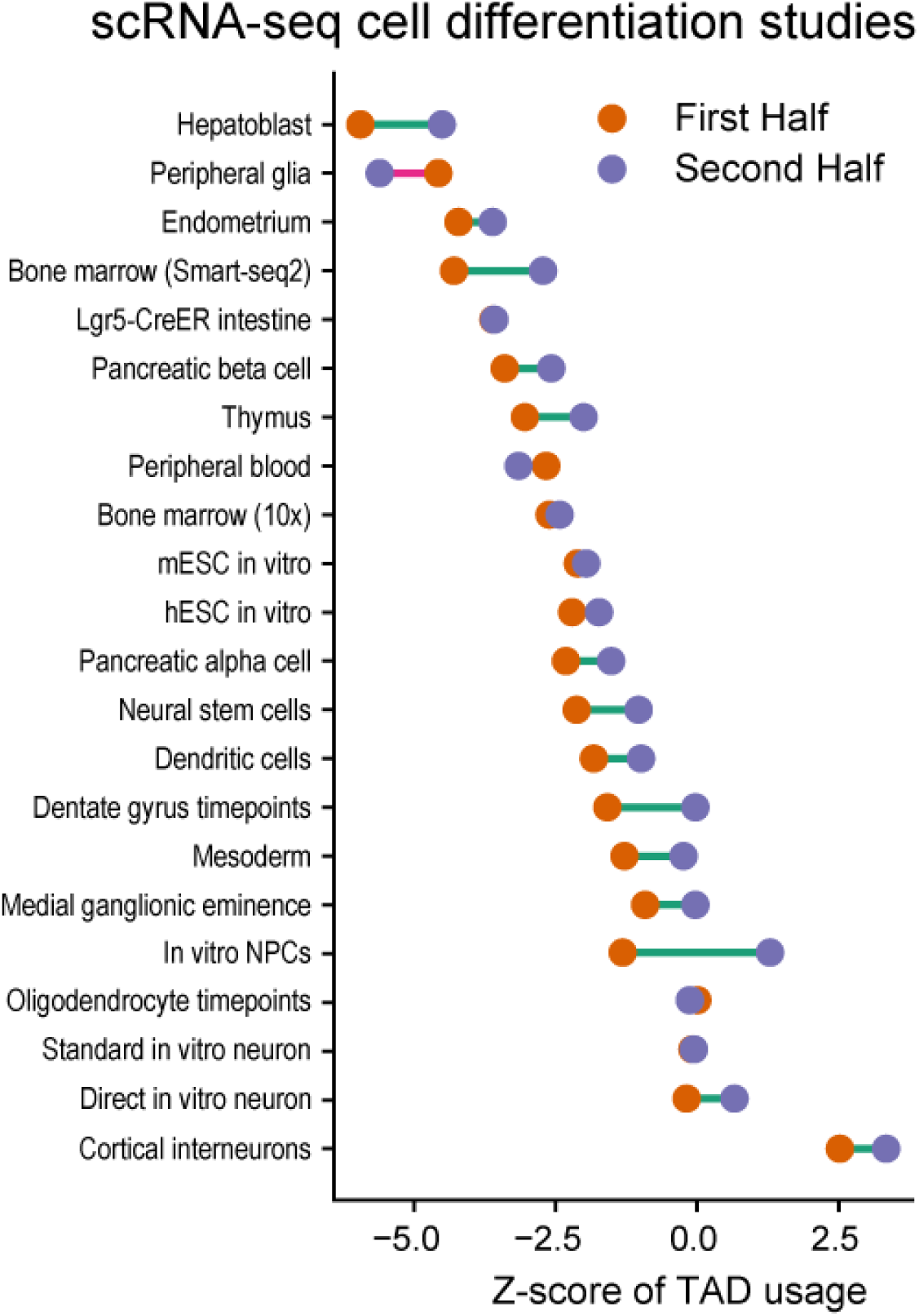
**Gene clustering into TADs decreases during cell differentiation**: This accompanies Figure 4D where we showed that cells earlier in the differentiation trajectory display greater clustering of genes into TADs. The partitioning of cells (per study) in that figure was done as “first stage vs the rest”, where the stage information was sourced from the *Order* variable in CytoTrace data [19]. In this figure, stages (i.e., *Orders*) are partitioned into two groups such that approximately equal number of cells are found in both groups. Regardless of the partitioning approach, the clustering observation holds.

**Figure S4:**
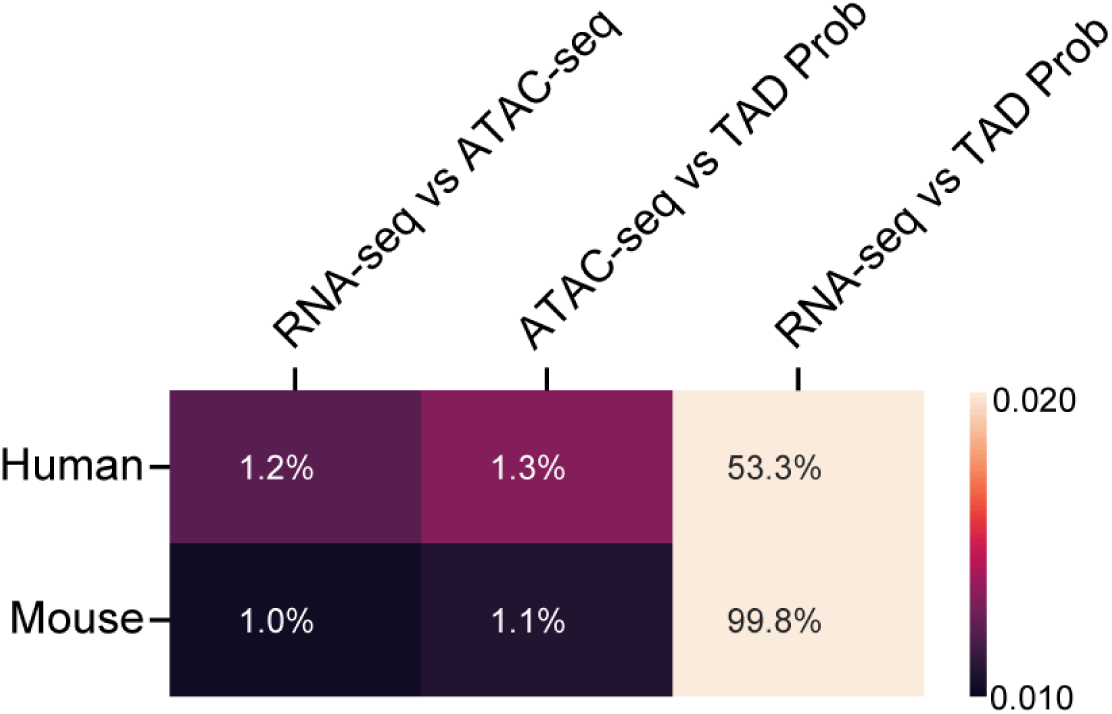
**TAD signatures improve agreement with ATAC-seq data**: This accompanies Figure 5A where we compared per-TAD activation probabilities and average RNA-seq scores against the presence of any ATAC-seq peaks. Here, we perform the same comparison with Spearman rank correlation against ATAC-seq peak counts as the measure. Again, TAD activation probabilities have a stronger correlation with ATAC-seq peak counts, though the sparsity of ATAC-seq results in very low correlations overall.

## Footnotes

1 We also define TAD’s “capacity” as the number of genes in its bag-of-genes representation. Thus, a TAD’s capacity is constant but its usage varies.

